# Integrative Spatial Omics for Systems-Level Mapping of Pathological Niches

**DOI:** 10.1101/2025.09.12.675904

**Authors:** Angela R.S. Kruse, Roy Lardenoije, Lukasz G. Migas, Claire F. Scott, Cody Marshall, Morad C. Malek, Adel Eskaros, Thai Pham, Kristie Aamodt, Madeline Colley, Lissa Ventura-Antunes, Melissa A. Farrow, Raf Van de Plas, Joana Goncalves, Matthew Schrag, Alvin C. Powers, Jeffrey M. Spraggins

## Abstract

Spatial ’omics technologies are a powerful tool for mapping the relationship between cellular organization and molecular distributions in healthy and diseased tissue microenvironments. Here, we describe a novel multimodal pipeline that represents experimental and computational advances for spatiomolecular analysis of tissue samples across molecular classes. This adaptable method integrates matrix-assisted laser desorption/ionization imaging mass spectrometry spatial lipidomics, spatial transcriptomics, protein imaging via multiplexed immunofluorescence microscopy, and histopathological staining to uncover spatiomolecular profiles associated with unique cellular niches and pathological features. We demonstrate the power of this approach using two different complex human disease systems: Alzheimer’s disease in human brain tissue and type 2 diabetes mellitus in the human pancreas. This work establishes and demonstrates a generalizable framework for multimodal spatial integration, enabling precise mapping of molecular mechanisms that underlie complex tissue pathologies.

## Introduction

Spatial biology is ideally suited to fully characterize complex and spatially heterogeneous tissue environments in healthy and diseased tissues. Spatial proteomics, spatial metabolomics, and spatial transcriptomics in particular have emerged as powerful tools for biological discovery [1–11]. While individual technologies are advancing rapidly, there is an urgent need for multimodal frameworks to enable comprehensive, systems-level analysis of spatial ‘omics data. Here, we present an adaptable workflow for the generation, co-registration, and cross-modality mining of spatial datasets to maximize biological insights from spatial ‘omics data.

The need for spatial analysis is clear in complex disease systems such as Alzheimer’s disease (AD) and type 2 diabetes mellitus (T2D), which represent ideal models to demonstrate the power of multimodal spatial analysis in the brain and pancreas. AD is the most common form of dementia, with an estimated 6 million adults in the United States diagnosed with the disease in 2020 [12]. Neuritic plaques, composed of extracellular β-amyloid deposits, are a characteristic neuropathological feature associated with AD [13]. Cerebral amyloid angiopathy (CAA) is an AD-associated pathology co-occurring in 82% to 98% of AD patients [14]. It is characterized by β-amyloid deposits forming within the walls of cerebral arterioles and occasionally capillaries [15–20]. T2D is a global health crisis impacting approximately 1 in 10 adults globally and leading to life-threatening complications and decreased life expectancy [21, 22]. T2D is associated with relative insulin deficiency due to peripheral insulin resistance and the dysfunction and loss of insulin-producing beta cells within pancreatic islets. Amyloid plaques formed from aggregated islet amyloid polypeptide (IAPP) have been observed at increased frequency in the islets of individuals with T2D [23, 24]. However, the relationship between these amyloid plaques and the molecular causes of beta-cell dysfunction and changes in islet composition in T2D remains unclear. Lipids play active roles in both AD and T2D, influencing protein aggregation, neuronal membrane dynamics, and cellular metabolism [25–30]. Mapping their spatial distribution provides a critical layer of molecular context that complements transcriptomic and proteomic analyses.

Most studies of molecular changes in the brain in AD and pancreas in T2D have been limited to bulk tissue samples, dispersed single cells, and/or spatial analyses that focus on one molecular class [25, 27, 28, 31, 32]. Subsequently, relatively little is known about the cellular and spatial context of these pathologies. To address this, we developed an approach to integrate spatial lipidomics via imaging mass spectrometry (IMS), spatial transcriptomics (ST), and highly multiplexed immunofluorescence (MxIF) along with histopathology to enable the cross-modality mining across spatial datasets.

Matrix-assisted laser desorption ionization (MALDI) imaging mass spectrometry (IMS) is used here to perform untargeted lipid IMS on sections of fresh-frozen human brain and pancreas tissue. IMS is a powerful untargeted spatial omics technique capable of producing heatmap-style ion images that display the distribution of each analyte across a tissue section [33–36]. MALDI IMS workflows enable the detection of multiple analyte classes, including peptides, proteins, N-linked glycans, small-molecule drugs, metabolites, and lipids [34, 35, 37, 38]. Since IMS is based in mass spectrometry, it allows for structural characterization of analytes with high chemical specificity [7]. In this case, hundreds of lipids can be detected in a single MALDI IMS experiment, making this an ideal technology to spatially interrogate dysregulated lipid metabolism in disease and inflammatory states.

While high spatial resolutions can be achieved with IMS (<10 µm pixel size) [7, 8, 39], single-cell resolution is not routinely achievable. Therefore, lipid IMS is complemented by more targeted single-cell-level spatial transcriptomics and proteomics. ST is an emerging technology in spatial biology [40]. Among the available approaches, imaging-based technologies generally enable the detection of transcripts from mesoscale to single-cell resolution. We demonstrate our integrative pipeline using the 10X Genomics Xenium platform, which utilizes circularizable DNA probes to target hundreds to thousands of transcripts *in situ* at subcellular resolution [41]. Signal enhancement is then performed using rolling circle amplification before cyclical hybridization of fluorescent-bound probes and imaging, leading to increased sensitivity of the assay in detecting low-abundant transcripts. After detection, transcripts are localized to cells based on cell boundaries inferred through nuclear segmentation, enabling single-cell-level analysis of tissue microenvironments.

The non-destructive nature of the ST workflow preserves tissue morphology for post-ST analyses such as microscopy-based protein imaging. As such, we combined ST *in situ* gene expression data with PhenoCycler MxIF on the same tissue section. This platform enables the multiplexed and cyclic detection of tens of proteins using antibodies conjugated with nucleic acid barcodes [42]. MxIF is a high-resolution targeted technology, a standard for cell type profiling and phenotyping, and ideal for mapping vascular elements. Since fluorescent reporters are removed after MxIF imaging, small-molecule staining can be performed on the same tissue sections.

Spatial ‘omics data are harmonized through image registration to enable accurate high-resolution integration and analysis. We demonstrate multiple supervised and unsupervised strategies for cross-modality mining. Although AD and T2D are distinct diseases, we apply this pipeline to compare amyloid pathology in the brain and pancreas, illustrating the power of multimodal integration across molecular classes and organ systems. Beyond the pathological models analyzed here, this workflow provides a broadly applicable approach for multimodal spatial analyses across diverse tissues and disease contexts. As spatial technologies continue to expand in scale and precision, such integrative strategies will be essential to translate molecular maps into actionable biological insight.

### Design [This section is supposed to highlight novelty in a technology article]

Multimodal spatial analysis has been constrained by the difficulty of connecting untargeted molecular data to specific cell types and functional or pathological tissue regions. Our pipeline overcomes this by integrating high-resolution, targeted single-cell technologies with untargeted imaging mass spectrometry to facilitate systems-level discovery across spatial and molecular class scales. This framework bridges chemically rich, spatially resolved IMS data with transcriptomic and proteomic context, enabling mechanistic insights into spatially defined disease processes such as amyloid pathology in AD and T2D.

## Results

### Multimodal Molecular Imaging

In this study, we used tissue serial sections from each of our disease systems: human AD brain and T2D pancreatic tissues. Autofluorescence microscopy was collected on all tissues before and/or after molecular imaging to enable spatial co-registration between serial tissue sections [43, 44]. On the first tissue section, we performed ST using a pre-designed brain panel and multi-organ tissue mapping panel for the brain and pancreas, respectively (Figure 1). After ST, we performed MxIF [42]. DAPI nuclear staining performed in both the ST and MxIF workflows was used for image registration between modalities. After MxIF, we performed staining for amyloid using thiazine red for the brain and thioflavin S for the pancreas. On the second serial tissue section, we performed spatial lipidomics using MALDI IMS at 10 µm pixel size (Figure 1). This approach, which utilizes only two serial tissue sections, enables the measurement of tens of proteins, hundreds of transcripts, and hundreds of lipids, as well as any histopathological markers of interest.

**Figure 1.**
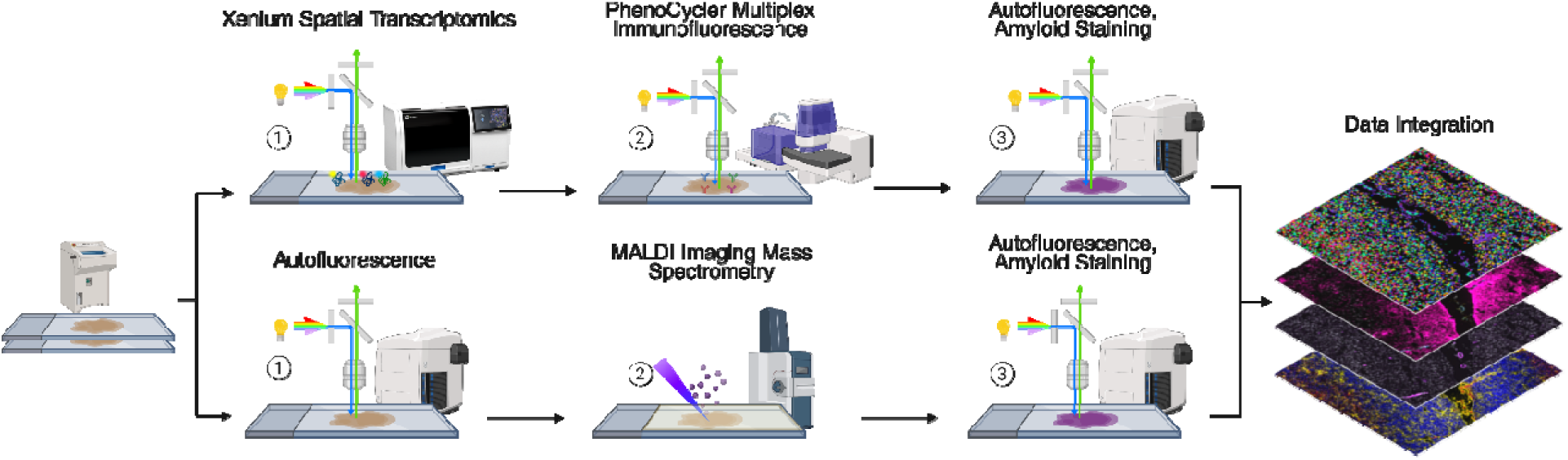
Multimodal analysis workflow. Two serial tissue sections were used in this workflow. On the first section (top), spatial transcriptomics (with Xenium) was performed, followed by spatial proteomics (with PhenoCycler multiplex immunofluorescence), autofluorescence microscopy, and amyloid staining. On the second section, autofluorescence microscopy was performed, followed by spatial lipidomics (with MALDI IMS) and post-IMS autofluorescence microscopy.

The pre-designed Xenium gene panels (Table S5) include markers for most major tissue structures and cell types. For instance, DCN and PECAM1 can be used as gene markers for vasculature in the brain (Figure 2A). In the pancreas, GCG and AMY2A successfully mark cell types within the islets and exocrine tissue, respectively (Figure 2E). The Xenium cell segmentation method relies on a nuclear stain, which can result in incomplete or inaccurate cell boundary markings. By including structural protein markers such as e-cadherin and GFAP in the MxIF panel, we were able to confirm cell boundaries and contiguous tissue structures, including vasculature (Figure 2B,F). Moreover, small-molecule stains for amyloid were used to mark amyloid-positive structures within the tissue for further downstream investigation (Figure 2C,G). On the serial section, lipid distributions were measured with MALDI IMS, which allows for the measurement of the relative abundance and localization of lipids in an untargeted manner in each tissue sample (Figure 2D,H). Peak detection resulted in 1,420 IMS peaks in the brain and 889 in the pancreas, including signals associated with matrix and isotopes. Peak annotation was performed via mass accuracy using a reference database of lipid masses [45, 46] and allowed for tentative annotation of 504 lipid peaks in the brain and 261 in the pancreas, each with unique spatial localizations (Figure S1, Table S1). For example, PA 36:1 was detected in the white matter of the brain, whereas a PaZ-PC lipid was detected in the vasculature within the meninges (Figure 2D). A SHexCer 42:1 lipid was detected within pancreatic islets, and its localization correlates most closely with glucagon-producing alpha cells identified via MxIF (Figure 2F,H). Ceramide phosphates were found to vary in localization based on chain length and degree of saturation in the pancreas; CerP 34:1;O2 was detected in islets and vascular tissue, whereas CerP 38:2;O2 was detected in acinar tissue (Figure 2H).

**Figure 2.**
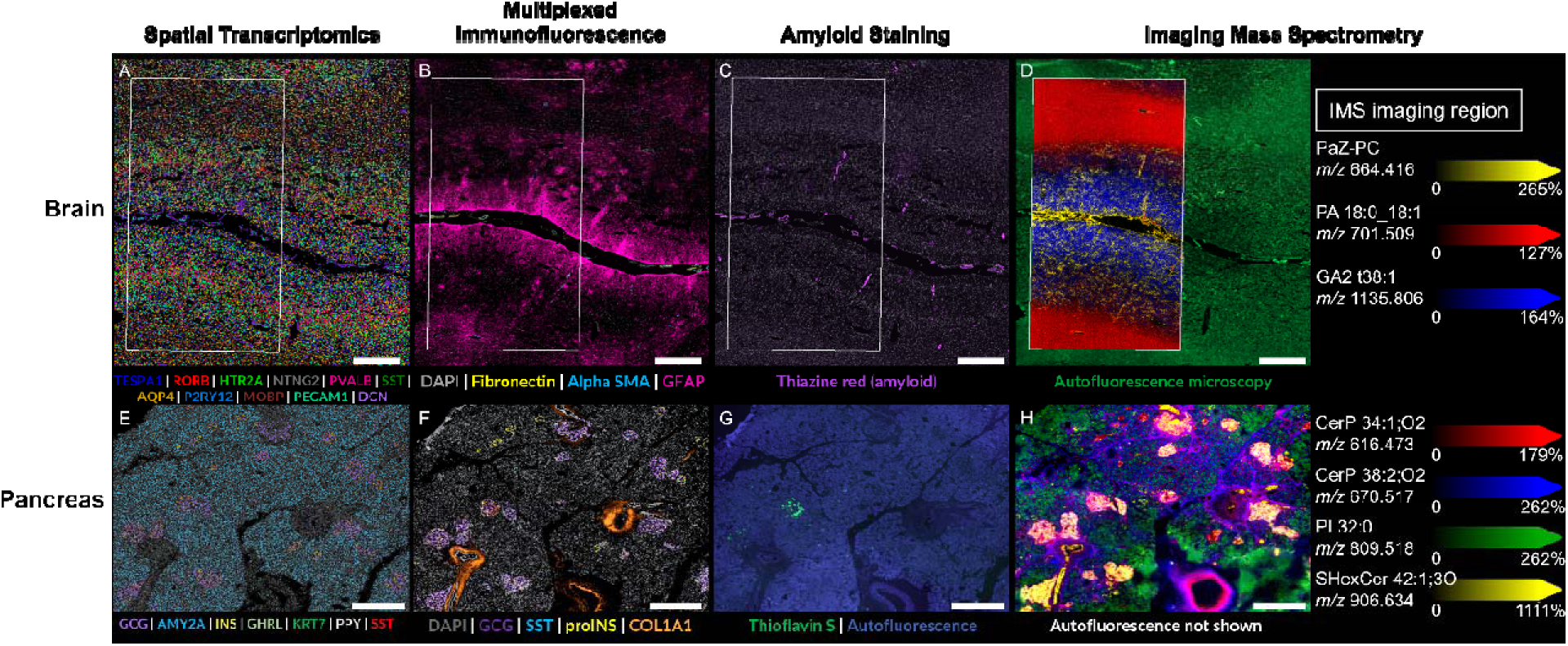
Multimodal imaging in brain tissue from a donor with severe Alzheimer’s disease (top row) and pancreas tissue from a donor with type 2 diabetes (bottom row). A,E) Spatial transcriptomics (ST; Xenium) was used to establish transcriptional profiles at sub-cellular resolution. Cell segmentation was performed using a DAPI nuclear stain and nuclear expansion. Cell overlay color indicates marker genes. B,F) After ST, multiplexed immunofluorescence (MxIF; PhenoCycler) was performed to facilitate annotation of cell types and vascular elements. C,G) Following MxIF, thiazine red (brain) or thioflavin S (pancreas) staining was performed to identify amyloid aggregates. D,H) On a serial tissue section, autofluorescence microscopy (green) was performed to facilitate image registration, and spatial lipidomics (MALDI Imaging Mass Spectrometry (IMS)) was performed to analyze the distribution and relative intensities of lipids in an untargeted manner. IMS was performed with a pixel size of 10 µm. Lipids were annotated via exact mass matching with the classes plasmalogen (PaZ-PC), phosphatidic acid (PA), ganglioside (GA), ceramide phosphate (CerP), phosphatidylinositol (PI), and sulfatide (SHexCer).

The spatial nature of this multimodal dataset allows us to apply multiple cross-modality mining approaches to gain biological insight (Figure 3). The crucial first step for any multimodal spatial biology study is accurate image co-registration. Since the various modalities were collected as part of separate experiments, often with varying pixel sizes, and some on serial sections, it was necessary to align each spatial datasets into a unified coordinate system. To achieve this, we utilized an in-house developed software, *image2image*, which serves as both a visualization and registration tool with a graphical user interface (GUI) for defining anchor points, enabling semi-automated registration, and assessing registration accuracy (Figure S2) [47–48]]. Following the completion of these registration steps, a visual inspection of all images was performed at both tissue and cellular scales to ensure accuracy (Figure S3).

**Figure 3.**
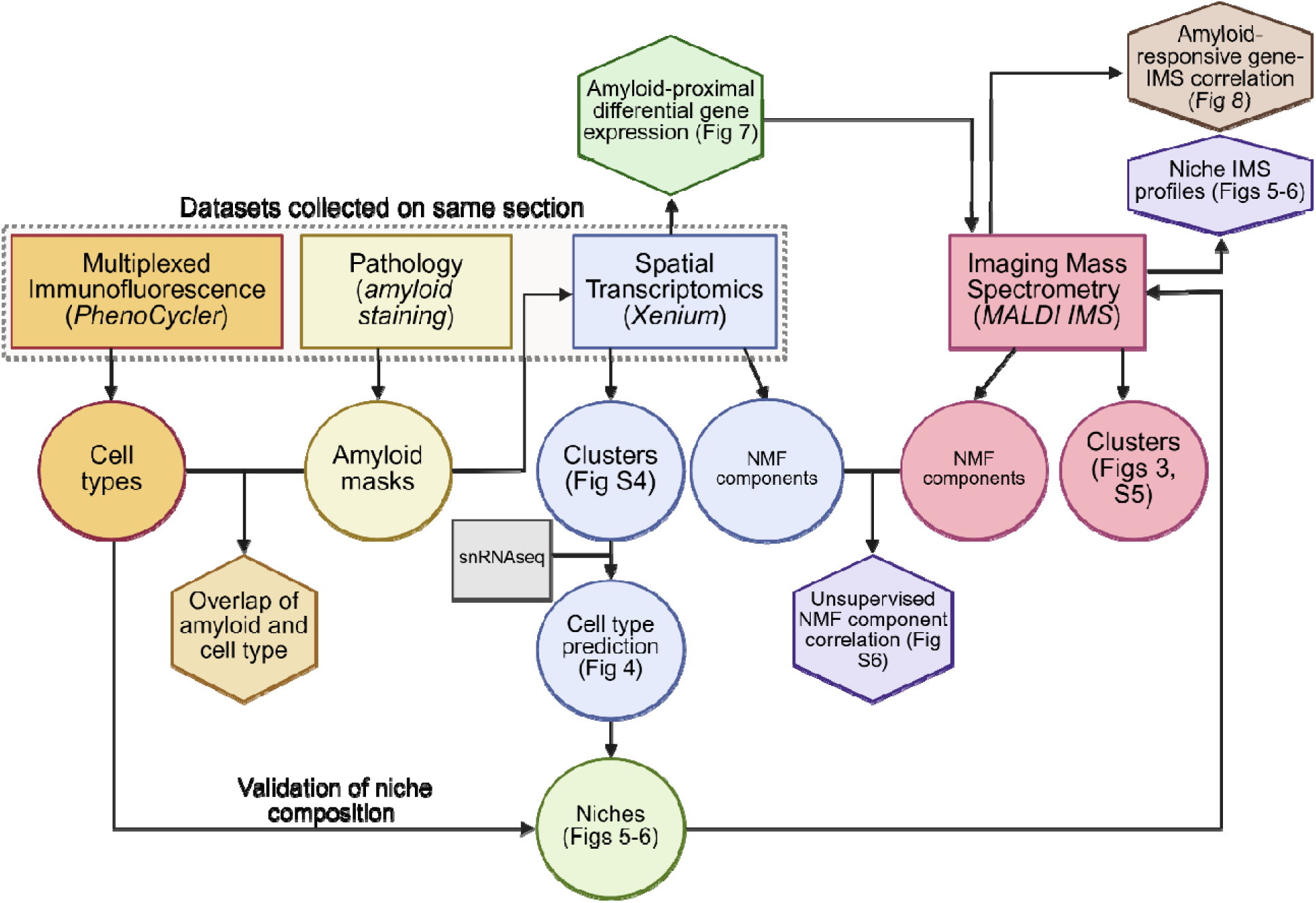
Cross-modality data mining workflow. Primary datasets (shown in rectangles) include MxIF, amyloid staining, and ST collected on the same tissue section (indicated with a dashed rectangle), and MALDI IMS collected on a serial section. Secondary analyses (shown in circles) include cell type information, amyloid segmentations, clusters from ST and IMS, and nonnegative matrix factorization (NMF) components from ST and IMS. These analyses are used for cross-modality mining (shown in hexagons), such as correlation of NMF components between datasets.

### Unsupervised Analysis

Unsupervised clustering is well suited for initial parsing of large, unlabeled datasets, such as those generated by omics platforms. Here, we applied unsupervised clustering separately to the ST (Figure S4) and IMS (Figures 4, S5) datasets to identify spatial regions with molecular similarities at the transcript and lipid levels, respectively. In the brain, the clustering of both ST and IMS resulted in the separation of the meninges, white matter, and gray matter, in addition to radially distinct regions with molecularly unique signatures (Figures S4C, 4A). Similarly, clustering of the pancreas data separated acinar, islet, vascular, and ductal tissues (Figures S4D, S5). Each IMS cluster is associated with an average mass spectrum (Table S2). These cluster-associated spectra in the brain are visibly distinct, and *m/z* values can be ranked by their association coefficient with each cluster (Figure 4B-I). Several ions, including *m/z* 572.277, 701.513, and 906.633, were associated with multiple clusters, emphasizing that the distinction of k-means clusters is multivariate, based on a combination of intensity values from many *m/z* values (Table S2). Clustering in the pancreas demonstrated a distinct intra-section separation at the IMS level, likely due to fat infiltration (Figure S5B-G). However, functional tissue units were successfully segmented in both the IMS and ST data (Figures S4-S5).

**Figure 4.**
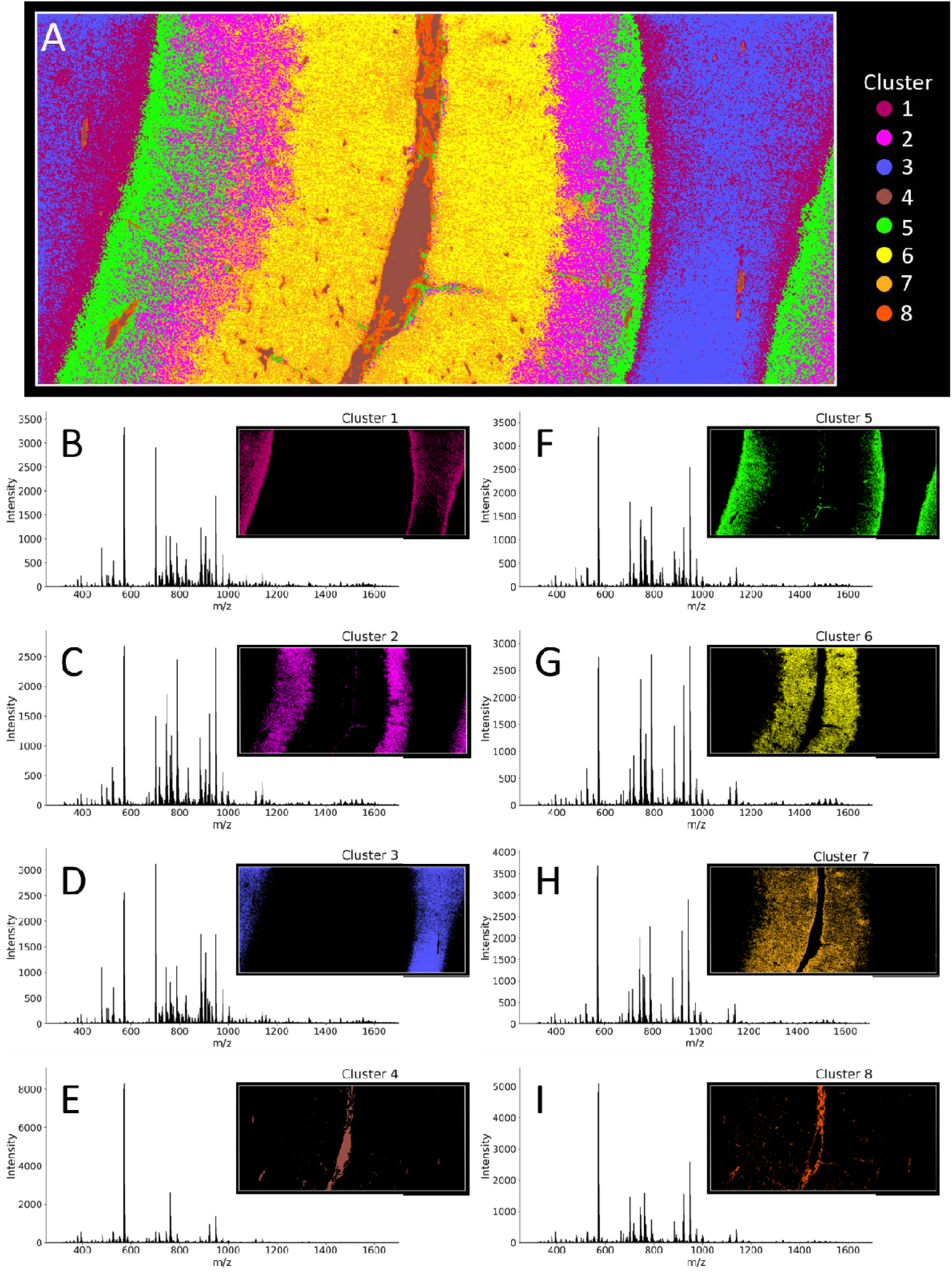
Unsupervised clustering of spatial lipidomics (MALDI IMS) data in AD-brain. K-means clustering was performed to segregate IMS data into 8 clusters. Cluster identities are spatially visualized as an overlay (A) and as individual cluster images associated with an average MALDI IMS spectral profile (B-I).

Using an orthogonal approach, dimensionality reduction via non-negative matrix factorization (NMF) was performed to parse ST and IMS data into interpretable components. Pearson’s correlation was performed to identify spatial similarities between ST and IMS components. In the pancreas, the greatest correlation was found between ST component 12 and IMS component 14. Both clusters contained molecules localized to islets (Figure S6). While clustering and NMF at the whole section level captured most gross anatomical features in both organs, there was, interestingly, no specific cluster that could be associated with the observed amyloid pathologies in either the brain or the pancreas. This is potentially due to the complex mixture of cell types associated with amyloid aggregates, particularly in the vasculature in the brain and islet tissue in the pancreas. Dimensionality reduction enabled a direct comparison of genes and lipids from the ST and IMS modalities, respectively (Figure S7).

Together, these unsupervised analyses effectively captured major tissue architectures and molecular gradients, highlighting broad spatial trends. However, these approaches are inherently limited in their ability to resolve smaller or biologically specific features, such as localized amyloid deposits or subtle cell-type boundaries. A targeted and supervised strategy is required to enable hypothesis driven interrogation of specific biological structures, pathologies, or cell-type distributions within spatial omics datasets.

### Cell Type Prediction

To facilitate targeted cross-modality analyses, cell type prediction using single-cell data was used to assign annotations to spatially segmented cells in ST data. We leveraged publicly available annotated reference datasets to perform cell type prediction in ST data from the brain (Figure 5A) and pancreas (Figure 5D) [49]. These predictions were then used to split the ST data and re-cluster it per cell type, leading to the identification of several cell type subclusters (Figure 5). Some of these subclusters showed clear regional distributions. For example, intratelencephalic (IT) cells showed clear spatial segregation in the brain (Figure 5B). Astrocytes predominantly segregated between the white and gray matter of the brain (Figure 5C). Alpha cells in the pancreas also demonstrate distinct clusters that appear to be associated with the inner islet compared to the islet periphery (Figure 5E). Pancreatic acinar cells appear to cluster based on proximity to the edge of the tissue section (Figure 5F).

**Figure 5.**
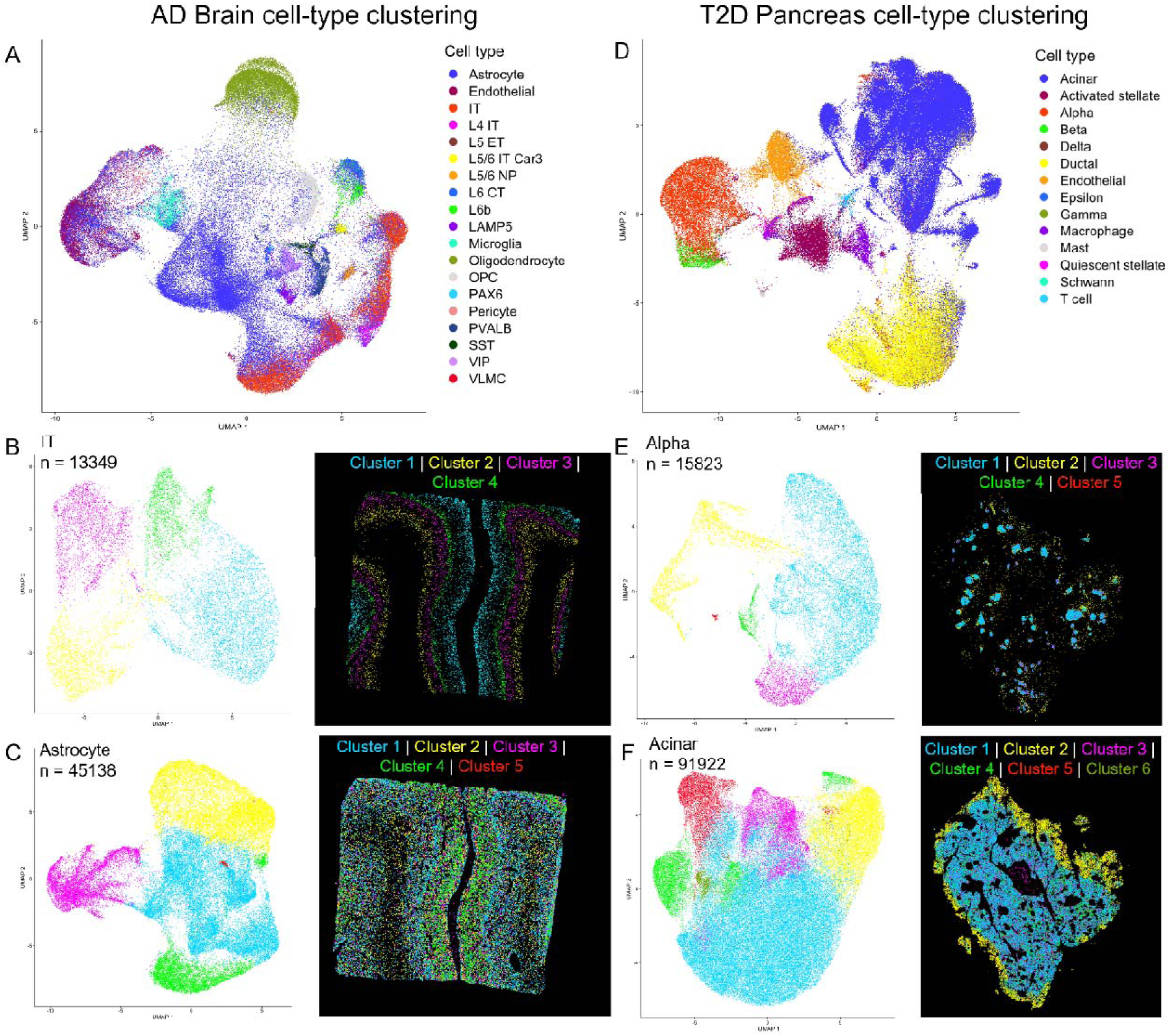
Cell type specific spatial transcriptomics (ST) data clustering. Cell type prediction was performed on the AD brain (left) and T2D pancreas (right) datasets. A. UMAP projections indicate predicted cell types in ST data from the AD brain. Predicted cell types include (from top to bottom of legend) astrocyte, endothelial, intratelencephalic (IT), layer 4 (L4) IT, layer 5 (L5) Extratelencephalic (ET), layer 5/6 IT Car3-positive (L5/6 IT Car3), layer 5/6 near-projecting glutamatergic neuron NP (L5/6 NP), layer 6 corticothalamic (L6 CT), layer 6b (L6b), Lysosomal-associated membrane protein family member 5 interneuron (LAMP5), microglia, oligodendrocyte, Oligodendrocyte progenitor cell (OPC), paired box 6 positive (PAX6), pericyte, parvalbumin positive (PVALB), somatostatin positive (SST), vasoactive intestinal polypeptide (VIP), Vascular leptomeningeal cells (VLMC). Additional clustering (left panels) and spatial visualization (right panels) were performed on specific cell types, including intratelencephalic (IT) cells (B) and astrocytes (C). D. Cell type predictions in T2D pancreas, including (from top to bottom of legend) acinar, activated stellate, alpha, beta, delta, ductal, endothelial, epsilon, gamma, macrophage, mast, quiescent stellate, Schwann, and T cell. Further clustering (left panels) and spatial visualization (right panels) were performed for alpha cells (E) and acinar cells (F).

### Niche Analysis

Using cell-type assignments informed by ST and snRNAseq data, a niche analysis was performed based on spatially co-occurring cell types. The cell type composition and spatial context of the different niches may provide some insight into their function (Figure 6A-B, 7A-D). For instance, in the brain, niche 6 encompasses the upper layers 1-3 of the cortex, characterized by the highest proportion of astrocytes, intratelencephalic neurons, as well as vasoactive intestinal peptide (VIP) expressing interneurons (Figure 6B). This is consistent with abundant Glial Fibrillary Acidic Protein (GFAP) staining from the SP data on the same tissue section (Figure S8). GFAP is a well-established astrocyte marker, and its proteomic localization offers an orthogonal confirmation of the high predicted astrocyte density within niche 6 [50]. Niche 1 represents a combination of cortical layers 4 and 5 due to the relatively high abundance of layer 4 intratelencephalic (L4 IT) and L5 extratelencephalic (ET) neurons (Figure 5B). Bordering niche 1 is niche 4, which has the most L5/6 IT Car3-positive and L5/6 near-projecting (NP) neurons. Niche 5 represents the deepest cortical layers, with the most L6 corticothalamic (CT) projection neurons and L6b neurons (Figure 6B). Niches 1, 4, and 5 are associated with the Na,K ATPase protein, which is expressed in epithelial cells and is involved in cell-cell adhesion (Figure S8F) [51]. White matter is primarily located in niche 3, which contains the most oligodendrocytes and oligodendrocyte precursor cells (OPCs). Niche 2 captures the vascular and meninges regions, as signified by the high proportion of endothelial cells, pericytes, and vascular leptomeningeal cells (VLMCs) (Figure 6A-B). This is orthogonally supported by the localization of the vascular-associated markers fibronectin and vimentin (Figure S8C-D).

**Figure 6.**
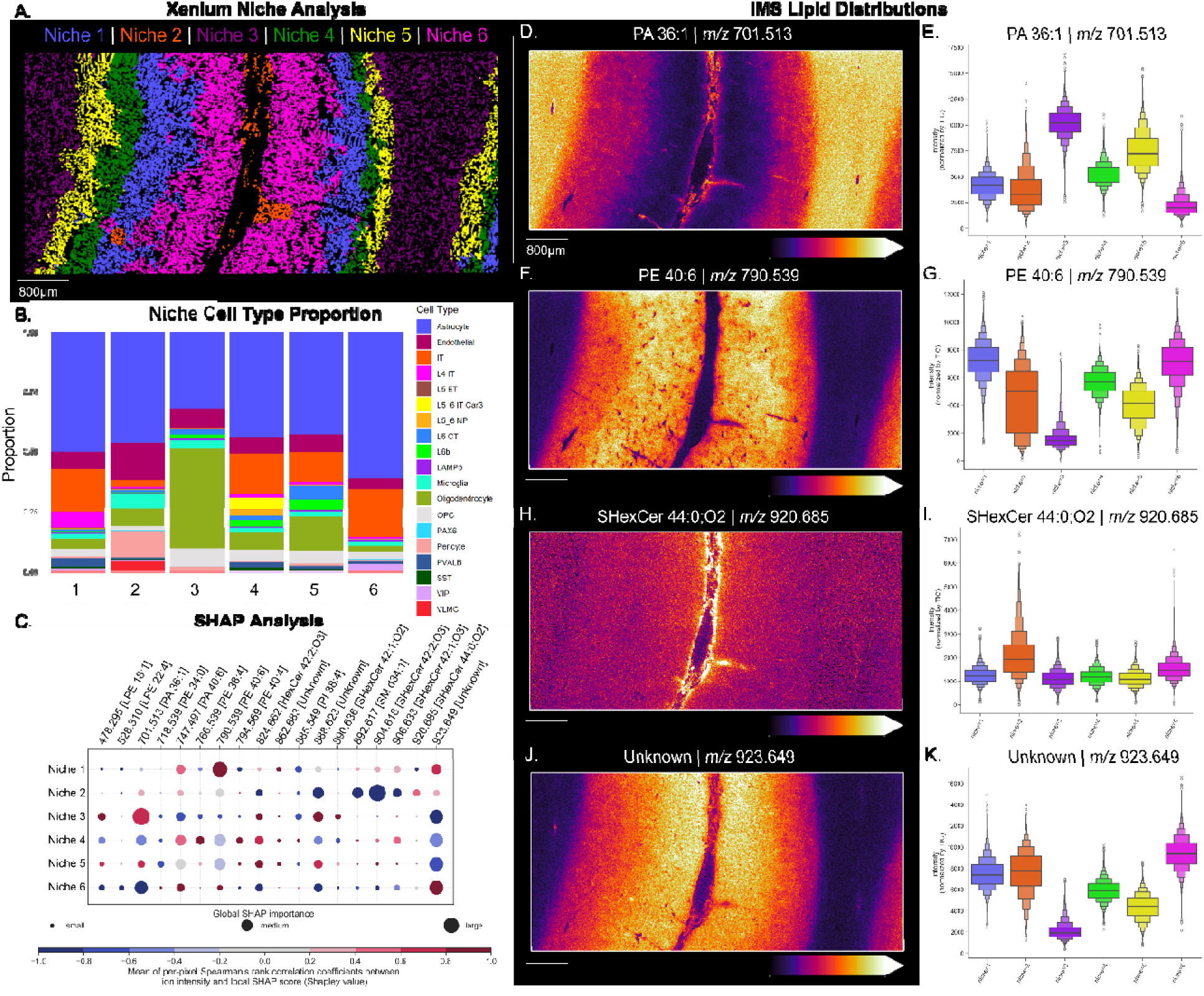
Cellular niche analysis in AD brain. A) Cellular niches predicted from spatial transcriptomics data. B) Cell type proportion within each niche. C) SHAP analysis showing lipids most predictive of each niche. D-K) Ion images and distributions for four ions with high SHAP importance scores and distinct spatial distribution within niches.

Five niches were predicted in the human pancreas (Figure 7A). Using MxIF imaging data as validation, we observe that niches 1 and 3 are within acinar pancreatic tissue, with niche 3 representing most of the acinar region, and niche 1 occupying small, circular regions that do not contain endocrine cells (Figure 7B-C). This distinction indicates exocrine tissue heterogeneity at the transcriptional level. Interestingly, while rare, all detected Schwann cells in the pancreas are allocated to niche 1, suggesting this may represent a neural niche within the exocrine pancreas (Figure 7D). Niche 2 (Figure 7D) appears to be a vascular and ductal niche, consistent with its representation of ductal and endothelial cells and spatial correlation with Col1A1 (Figure 7B-C). This niche also has relatively high proportions of immune cells, such as macrophages and T cells. Niches 4 and 5 are both islet niches, with niche 4 containing a higher proportion of beta and delta cells and niche 5 being dominated by alpha cells (Figure 7B-D).

**Figure 7.**
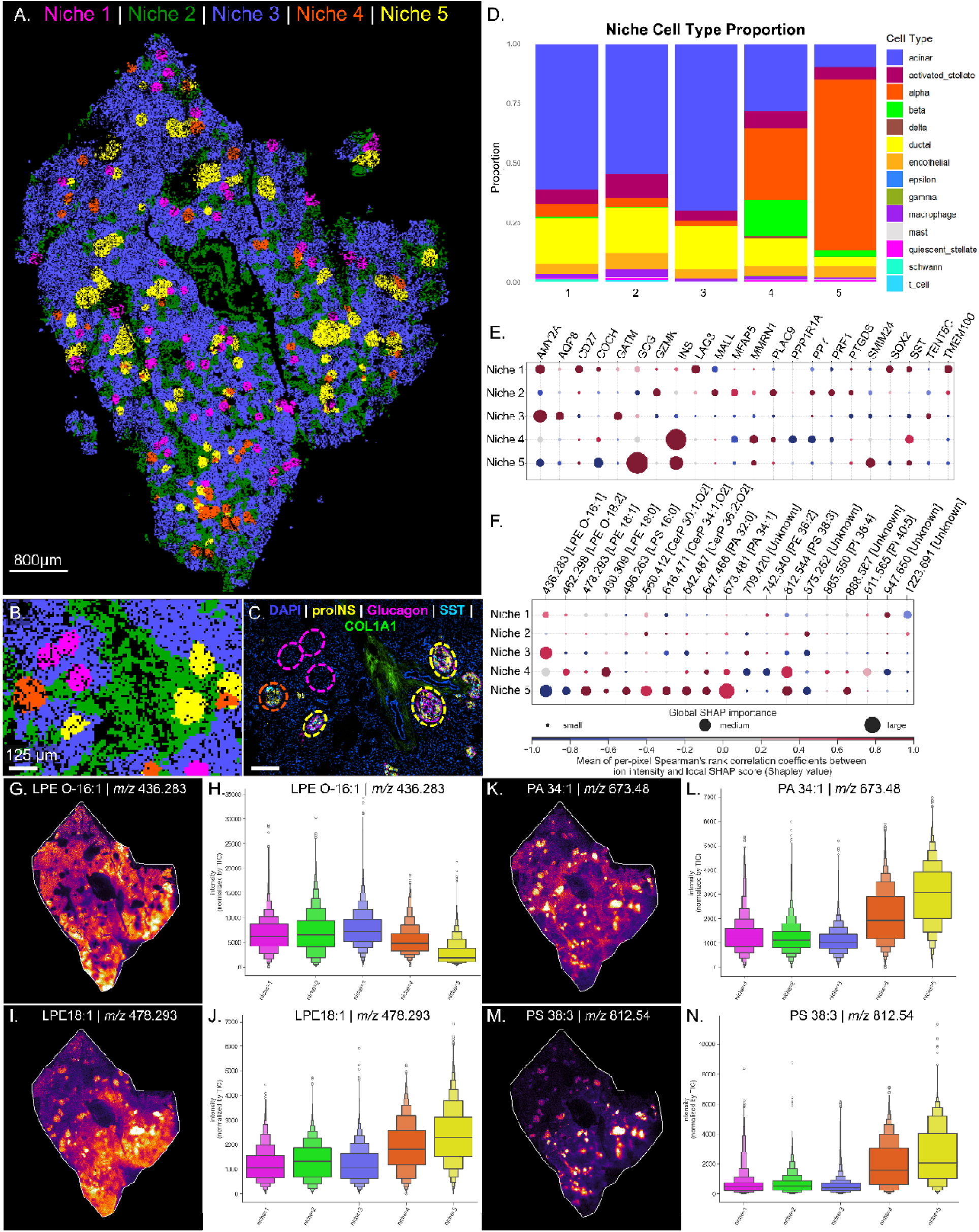
Cellular niche analysis in T2D pancreas. A-B) Cellular niches predicted from spatial transcriptomics data. C) Multiplexed immunofluorescence spatial proteomics showing key markers. D) Cell type proportion within each niche. E-F) SHAP analysis showing genes and lipids most predictive of each niche. G-N) Ion images and distributions for four ions with high SHAP values.

### Interpretable machine learning for biomarker discovery

The application of supervised interpretable machine learning allows for the identification of potential gene and lipid markers (*i.e.,* spatial biomarkers) from complex spatial datasets. Spatial niches determined from ST data were converted into binary masks and used as labels to train a classifier model via interpretable machine learning using Shapley additive explanations (SHAP) [52]. This approach enables unbiased identification of genes and lipids that contribute most significantly to distinguishing each spatial niche. To further interpret these results, feature abundance (*e.g.,* ion intensity, gene count) was correlated with local SHAP values using Spearman’s rank correlation. A positive Spearman correlation (shown in red in Figures 6C and 7E–F) indicates that higher signal intensity is associated with higher SHAP values, suggesting that the presence of the feature supports classification into the niche. Conversely, a negative correlation (shown in blue) indicates that lower signal intensity is associated with higher SHAP values, implying that the absence of the feature contributes to classification.

When applied to the ST data in the brain, we observe that Related Orphan Receptor B (RORB) and parvalbumin (PVALB) are positively associated with niche 1 (Figure S9). RORB is a marker for excitatory neurons in the entorhinal cortex, which are especially vulnerable in patients with AD [54]. PVALB is found in inhibitory interneurons [55]. Nephronectin (NPNT) was positively associated with niche 2, which contains a high proportion of endothelial cells. Myelin-Associated Oligodendrocytic Basic Protein (MOBP) was negatively associated with niches 2 and 6 but was positively associated with niche 3, containing a high proportion of oligodendrocytes (Figure 6B). Heparan Sulfate-Glucosamine 3-Sulfotransferase 2 (HS3ST2) and Heparan Sulfate-Glucosamine 3-Sulfotransferase 4 (HS3ST4) were positively associated with niches 4 and 5, respectively. Cut Like Homeobox 2 (CUX2) was positively associated with niche 6.

SHAP analysis was also applied to ST data within the pancreas. Amylase Alpha 2A (AMY2A) was predictive of both niches 1 and 3, which contain acinar cells. However, CD27, Lymphocyte-activation gene 3 (LAG3), SRY-box 2 (SOX2), and Transmembrane Protein 100 (TMEM100) were more highly associated with niche 1. Aquaporin 8 (AQP8) and glycine amidinotransferase (GATM) were more important for the classification of niche 3 (Figure 7E). As expected, the presence of insulin (INS) and somatostatin (SST) was important for the classification of niche 4, which contains the highest proportion of insulin-producing beta cells and somatostatin-producing delta cells, and the presence of glucagon (GCG) was important to classify the alpha cell-rich niche 5.

The use of niches to mine IMS data using interpretable machine learning is an especially intriguing approach, in that it allow for the correlation of transcriptionally unique multicellular regions to untargeted molecular IMS profiles for biomarker discovery. To demonstrate this, we applied interpretable machine learning to identify potential lipid biomarkers for each niche in the brain and pancreas (Figures 6C, 7F, S10, S11). In the brain, the presence of phosphatidic acid (PA) 36:1 was predictive of niches 3 and 5 (Figure 6C-E), phosphatidylethanolamine (PE) 40:6 was highly abundant in niches 1 and 6 but a stronger candidate biomarker for niche 6 (Figures 6C and 6F-G), and the sulfatide SHexCer 44:0;O2 was only a positive candidate biomarker for niche 2, and is most abundant that niche (Figures 6C and 6H-I). An unknown lipid was a strong candidate biomarker for niche 6, and correspondingly has high localization and abundance in that spatial region (Figure 6C and 6J-K).

In the pancreas, lysophosphatidylethanolamine (LPE) O-16:1 was found to be a marker for niche 3 (Figure 7F), and was more abundant in that niche compared to others (Figure 7G-H). This niche is dominated by acinar tissue, which is supported by the finding that AMY2A is a positive marker for this niche (Figures 7E-H). Phosphatidylserine (PS) 38:3 was a strong positive marker for niche 5 which is dominated by alpha cells, and a moderate marker for the classification of niche 4 which is dominated by beta cells (Figure 7F). This lipid was abundant in both islet niches, but most abundant in niche 5 (Figure 7M-N). LPE 18:1, PA 34:1, and lysophosphatidylserine (LPS) 16:0 were important for the classification in niche 5, which is dominated by alpha cells (Figures 7F, 7I-J, 7K-L). Interestingly, LPE 18:0 was most associated with niche 4, which is dominated by beta cells, indicating that a difference in one double bond allows for different localization within islets (Figure 7F). These niches are more biologically and spatially informed compared to the clusters, and as such form more discrete spatial neighborhoods. Niches are thus well-suited for integration with IMS, and niche-specific lipid profiles were determined for both organs (Figures 6-7).

### Molecular cartography of distinct histopathological features

The shared coordinate system of co-registered datasets allows for a topological approach to cross-modality data mining. This process can be applied to any unique histological feature identified in any of the modalities. To demonstrate this, in each of our disease models we determined the degree to which each cell co-localized, or overlapped, with an amyloid plaque. More specifically, amyloid plaque segmentations were performed using the small molecule stains thiazine red in the brain and thioflavin S in the pancreas. Then, using the shared coordinate system of the co-registered images, we observed the proximity of each cell (segmented from ST and MxIF images) to amyloid pathology. Additionally, we determined the distance from each cell to the nearest amyloid deposit (Figure 8A). Using this information to inspect subclusters within cell types, inherent differences within each cell type still prevailed over a potential amyloid-driven signal, as no clear subclusters associated with amyloid pathology were identified using this unsupervised approach.

**Figure 8.**
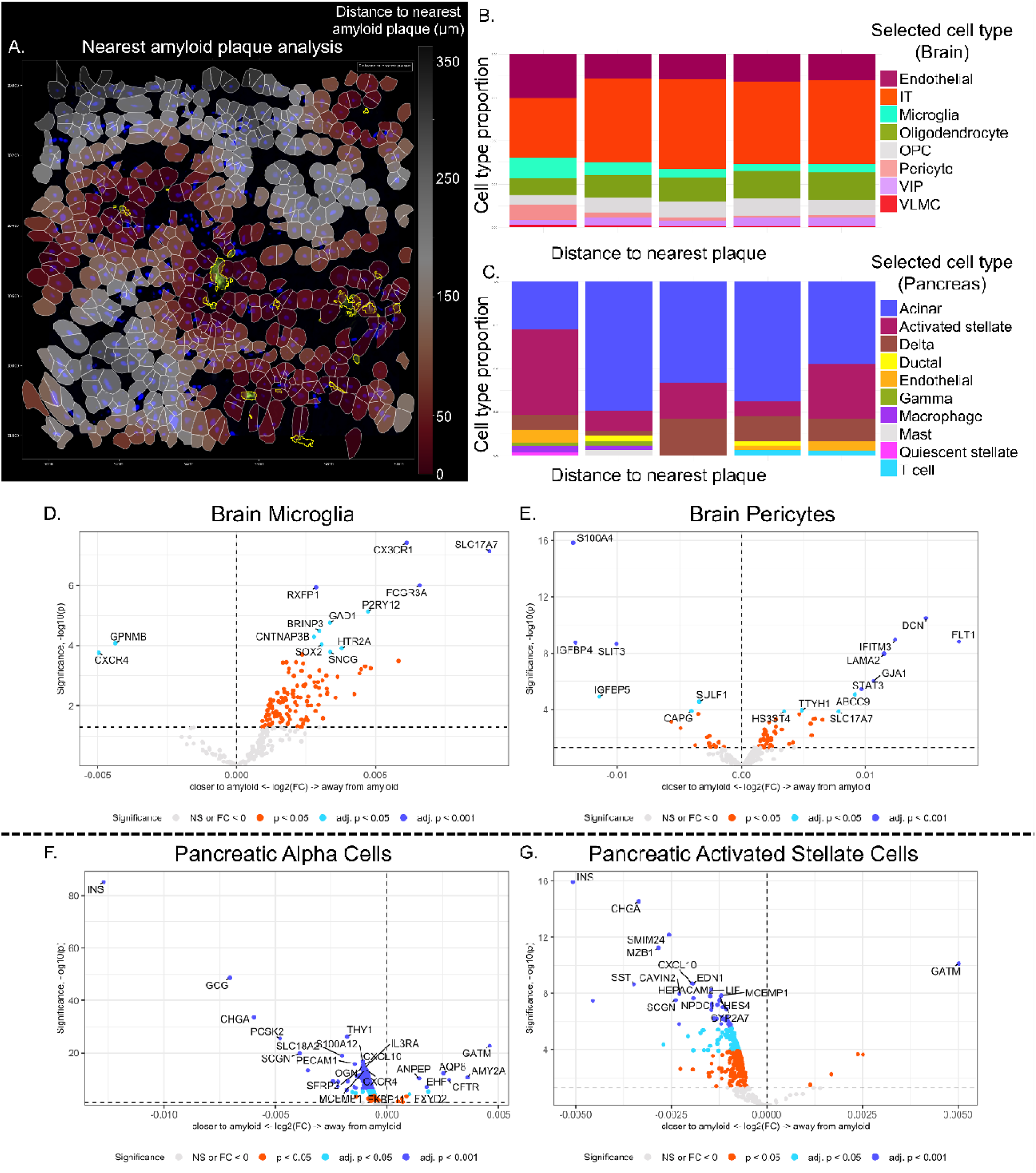
Differential gene expression in cell types based on proximity to the nearest amyloid plaque. A) Integrated view of the spatial transcriptomics cell segmentation (white border), nuclear DAPI staining (blue) and Thiazine Red staining (yellow) of an AD donor brain section. The segmented cells are colored (red gradient) by their distance to the nearest amyloid deposit. B) Cell type composition as distance to nearest plaque increases in AD brain. Cells are grouped into five categories: <10, 10-20, 20-30, 30-40, and 40-50 µm from the nearest plaque from left to right. C) Cell type composition as distance to nearest plaque increases in T2D pancreas. Cells are grouped into <10, 10-20, 20-30, 30-40, and 40-50 µm from the nearest plaque from left to right. D-G) Differential gene expression of cells within 100 µm of an amyloid aggregate comparing cells closest to amyloid to those further from amyloid AD brain microglia and pericytes (D and E, respectively) and T2D pancreatic alpha cells and activated stellate cells (F and G, respectively).

Using this molecular cartography approach, we begin by evaluating the changes in cell type proportion based on the distance from the nearest amyloid deposit (Figure S12, Figure 8). In the brain, the proportion of microglia is largest within 10 microns of an amyloid deposit (Figures S12A, 8B). The proportion of endothelial cells, pericytes, and VLMCs is also higher near amyloid plaques (Figure 8B). This is likely due to CAA, which is associated with amyloid aggregation within the vasculature. These proportions were not normalized to 100% to demonstrate that the total cell number decreases as the distance from amyloid plaques increases, since most amyloid plaques were found within vascular tissue. A similar confounding effect can be observed in the pancreas, where amyloid deposition is mainly localized to the islets, so islet-related cell types are present in higher proportions closer to amyloid (Figure S12B). To combat this, only cells in cluster 4 were selected, which primarily captures islets (Figure S4), to see if intra-islet changes in cell type proportions relative to amyloid could be observed. This approach revealed that macrophages and activated stellate cells have a slightly higher proportion closest to amyloid plaques in the pancreas (Figure 8C).

Given the representation of specific cell types near amyloid plaques in these different disease models, we performed differential gene expression analysis for each predicted cell type, comparing cells within 100 μm of an amyloid plaque to those further than 100 μm from a plaque (Figure 8D-G). Multiple cell types were found to have altered gene expression profiles based on their position relative to amyloid pathology. This included, most notably, astrocytes, microglia, endothelial cells, and pericytes in the brain, as well as acinar, alpha, and activated stellate cells in the pancreas (Table S3). In microglial cells within the brain, C-X-C Motif Chemokine Receptor 4 (CXCR4) and Glycoprotein Nmb (GPNMB) were up-regulated within 100 μm of amyloid (Figure 8D). Both mediate inflammatory responses, and GPNMB is associated with amyloidosis [56–58]. Solute Carrier Family 17 Member 7 (SLC17A7) and C-X3-C Motif Chemokine Receptor 1 (CX3CR1) were down-regulated in brain microglia near amyloid. SLC17A7 is associated with neuron-rich regions in the brain and is involved in glutamate transport. CX3CR1 is expressed in microglia and plays a role in amyloid deposition in the brain, though its role appears to be dose-dependent and variable based on the animal model studied [59, 60]. In pericytes, S100 Calcium Binding Protein A4 (S100A4), Slit Guidance Ligand 3 (SLIT3), Insulin Like Growth Factor Binding Protein 4 (IGFBP4), and Insulin Like Growth Factor Binding Protein 5 (IGFBP5) were up-regulated near amyloid, while Fms Related Receptor Tyrosine Kinase 1 (FLT1) and Decorin (DCN) were down-regulated (Figure 7E). S100A4 and SLIT3 are involved in cellular motility [61–63]. IGFBP4 and IGFBP5 are involved in insulin-like growth factor regulation, which impacts many cellular processes, including smooth muscle cell migration and have been implicated in neuronal injury recovery [64–66]. DCN plays a role in collagen fibril assembly [67].

In pancreatic alpha cells, glucagon (GCG) was up-regulated near amyloid plaques (Figure 8F). Insulin (INS) was also found to be up-regulated in alpha and activated stellate cells (Figure 8F-G), but this most likely indicates that cell segmentation captured some overlap of cell types, particularly within islets. Chromogranin A (CHGA), Proprotein Convertase Subtilisin/Kexin Type 2 (PCSK2), Solute Carrier Family 18 Member A2 (SLC18A2), Thy-1 Cell Surface Antigen (Thy1), S100 Calcium Binding Protein A12 (S100A12), Secretagogin (SCGN), and CXR4 were up-regulated in alpha cells near amyloid plaques (Figure 8F). CHGA and PCSK2 serve neuroendocrine functions [68, 69], and SLC18A2 functions in neurotransmitter transport [70, 71]. Thy1 is particularly important in neural cells in multiple organs, supporting a potential role for neural cells in amyloid pathology in the pancreas [72]. S100A12 is a calcium binding protein and acts as a pro-inflammatory protein [73]. Glycine Amidinotransferase (GATM), CF Transmembrane Conductance Regulator (CFTR), and Aquaporin 8 (AQP8) were down-regulated near amyloid (Figure 8F). CFTR and AQP8 are both involved in ion transport within pancreatic cells [74, 75]. In activated stellate cells, Marginal Zone B, B1 Cell Specific Protein (MZB1), C-X-C Motif Chemokine Ligand 10 (CXCL10), and Caveolae Associated Protein 2 (CAVIN2) were up-regulated, whilst GATM was down-regulated near amyloid plaques (Figure 8G). CXCL10 is involved in calcium storage, immune response, and macrophage polarization [76]. CAVIN2 binds to phospholipids and is involved in membrane remodeling [77, 78].

### Inter-organ analysis of the histopathological microenvironment

The identification of genes differentially expressed in proximity to distinct histological features, such as amyloid plaques, provides a unique opportunity to demonstrate the ability to mine IMS data and establish gene-lipid relationships within spatially distinct microenvironments using our multimodal method. In our case, we demonstrated this using the amyloid microenvironment. To establish gene-lipid profiles in the AD brain and T2D pancreas, we calculated the spatial correlation between the genes found to be differentially expressed near amyloid plaques and ions detected with IMS (Figure 9, Figures S13-14). This analysis was performed using differentially expressed gene density for all segmented cell types in the brain and pancreas (Figures S13-14). Since IMS was performed on a serial section, this whole-sample approach is less susceptible to variation between sections than an analysis using single-cell segmentations. In the brain, an unannotated lipid at *m/z* 524.279 was spatially correlated with expression of the gene SLC17A7 (Figure 9A-B). Several lipids, including LPE 18:1 at *m/z* 478.295, were spatially correlated with MOBP in the brain (Figure 9C-D, S13). PA 36:1 at *m/z* 701.509 was correlated with the expression of FBLN1 (Figure 9E-F). In the pancreas, PA 34:1 at *m/z* 673.481 was correlated with SMIM24 (Figure 9G-H). PA 36:1 at *m/z* 701.513 was spatially correlated with FBLN1 in the pancreas as well as the brain (Figure 9I-J).

**Figure 9.**
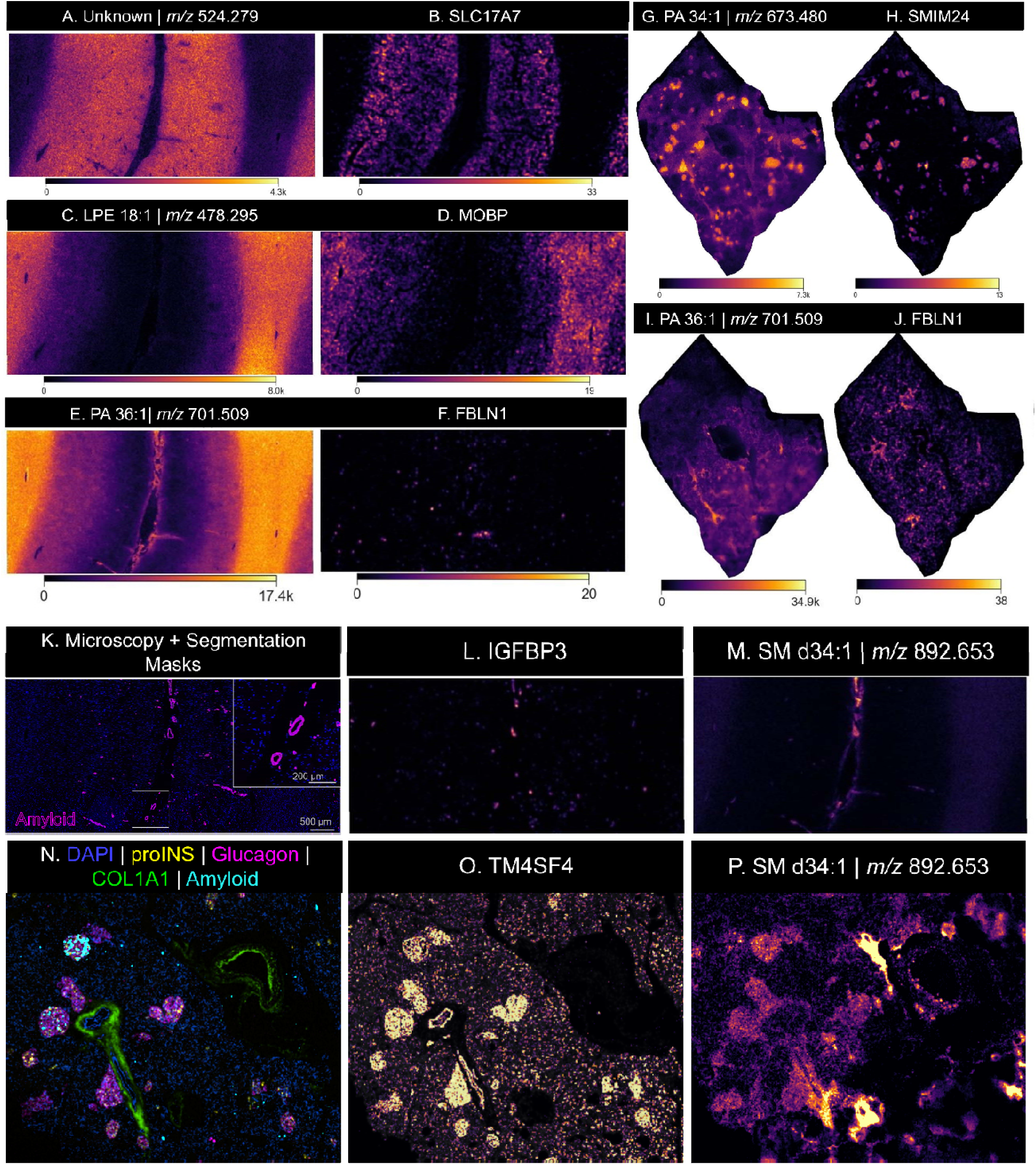
Spatial correlation of amyloid-responsive genes and lipids detected with IMS. A,C,E) show IMS ion images of *m/z’s* 524.279, 478.295 (LPE 18:1), and 701.509 (PA 36:1), respectively, in the brain. B,D,F) show gene density plots of SLC17A7, MOBP, and FBLN1, respectively, in the brain. G,I) show images of *m/z* 673.480 (PA 34:1), and 701.509 (PA 36:1) in the pancreas. H,J) show gene density plots of SMIM24, and FBLN1. K) DAPI and thiazine red staining to identify nuclei (blue) and amyloid plaques (magenta) in AD brain, respectively. Amyloid segmentations are shown in magenta. L) Gene density plot for IGFBP3 in the brain. M) Ion image of *m/z* 892.653 (SM d34:1) in the brain. N) Co-registered multiplexed immunofluorescence microscopy (MxIF) and thioflavin S amyloid staining in T2D pancreas. Selected MxIF channels proinsulin (proINS, yellow), glucagon (magenta), and collagen 1A1 (COL1A1, green) are displayed for visualization of islets and vasculature. Amyloid plaques stained via thioflavin S are shown in cyan. O) Gene density plot for TM4SF4 in the pancreas. P) Ion image of *m/z* 892.653 (SM d34:1) in the pancreas.

Furthermore, our multimodal method enables cross-organ comparisons between select histological features. Although the protein aggregates and pathophysiology of amyloid deposition differ in AD and T2D, we utilized amyloid deposits in AD brains and T2D pancreas to demonstrate the ability for cross-organ comparison of histological features with our model. To identify commonalities in transcriptional responses to amyloid proximity between the brain and pancreas, the differentially expressed genes across all cell types were compared between the two organs. A total of 16 genes were associated with amyloid proximity in both the brain and pancreas (Table S4). One such gene, FBLN1, was spatially correlated with PA 36:1 at *m/z* 701.513 in both the brain and pancreas (Figure 9E-F, I-J). FBLN1 is associated with fibronectin within the extracellular matrix. Interestingly, FBLN1 was down-regulated near amyloid in the brain and up-regulated in the pancreas, suggesting nuanced and organ-specific responses to amyloid pathology in these specific diseases. Additional genes, including EGFR, SOX2, CXCR4, CAV1, and CCL5, were amyloid-responsive in both organs (Table S4). The lipid SM d34:1 was found to be spatially correlated with amyloid-responsive genes in the brain and pancreas and is co-localized with amyloid pathology stains (Figure 9K-P). In the brain, genes including IGFBP3 share a spatial localization with segmented amyloid plaques and SM d34:1 (Figure 9K-M). In the pancreas, SM d34:1 is correlated with other amyloid-responsive genes, including TM4SF4 (Figure 9N-P). This lipid was detected within the vasculature in the brain but was found in pancreatic islets and vascular tissue (Figure 9M,P). The global approach linking amyloid-responsive genes to lipids demonstrates how this technique provides a unique opportunity to link these types of profiles across molecular classes and organ systems.

## Discussion

In this study, we introduce a novel multimodal approach to map complex tissue environments across molecular classes in health and disease. By integrating mass spectrometry with targeted single-cell ST, MxIF, and pathology staining, we generated a chemically rich and biologically informative dataset. This method increases the utility of untargeted lipid profiling by contextualizing it with orthogonal spatial modalities. Previous work demonstrates the power of integrating IMS and microscopy [39], and here we expand this workflow to ST and histopathological small-molecule stains. Our approach is broadly adaptable to other IMS modalities and fluorescence-based spatial omics platforms, offering a generalizable framework for multimodal data integration (Figure 3).

A key strength of our methodology lies in the combination of image co-registration, spatial segmentation, unsupervised and supervised analytical approaches, and the conscientious integration of datasets with differing spatial resolutions. Image co-registration was performed using an in-house developed software, *image2image*, which serves as both a visualization and registration tool [47]. Using *image2image*, each dataset from both the same tissue section and serial tissue sections was integrated into a common coordinate system. Segmentation of tissue regions and features was performed using unsupervised clustering, niche analysis, and cell type segmentation. This process is flexible and can be tailored to each researcher and their specific scientific question. The integration of same-section and consecutive-section data allowed for a comprehensive multi-scale analysis, enabling single-cell resolution data integration where possible.

Previous work has co-analyzed ST and IMS as well as MxIF and IMS, but this is the first approach to our knowledge that integrates IMS, MxIF, ST, and small-molecule pathology stains. To illustrate the effectiveness of our approach, we performed intra-organ analysis of human AD brain and human T2D pancreas tissues in addition to demonstrating how this method can also be used for cross-organ comparison, using the feature of amyloid plaques to demonstrate this type of analysis. Our approach is especially suited to investigating the role of lipid metabolism in these disease systems, which have spatially complex pathophysiology in affected tissues.

Our findings in the AD brain and T2D pancreas were consistent with prior studies that have investigated single molecular classes, in addition to providing new insights from cross-modal comparisons. For instance, previous studies have shown that phosphatidylcholine, phosphatidylethanolamine, and phosphatidylinositol levels are significantly decreased in the neuronal membranes of AD patients [32, 130–134]. In the brain, we observe that PE 40:6 is strongly associated with niche 1, which is enriched in astrocytes and IT cells (Figures 6B-C). In the pancreas, LPE lipids including LPE O-16:1, LPE O-18:2, LPE 18:1. And LPE 18:0 were important for the classification of acinar compared to islet niches. PE 36:2 and PI 38:4 were also assigned high SHAP importance values to classify pancreatic niche 4, which is dominated by beta cells. We also found that CAVIN2, which encodes a phospholipid-binding protein, is up-regulated near amyloid plaques in pancreatic activated stellate cells (Figure 8G). Future studies may further evaluate the spatial role of phospholipids and their regulatory gene partners in these two diseases.

Several lipids, including sphingolipids, have been observed to change in abundance and localization in the AD brain as well as in a macaque Parkinson’s disease model [79–81]. Our observation of SM d34:1 in proximity to amyloid plaques in the brain and pancreas suggests that conserved sphingomyelin lipids may be involved in plaque formation in AD and T2D (Figures 9Q and 9T). This is supported by known sphingomyelin dysregulation during amyloid aggregation in the brain [82]. Our approach demonstrates robust detection of lipid species relevant to both AD and T2D. Future work can apply this to additional samples to assess the relevance of each of these lipid molecules during disease progression.

We observed increased proportions of microglia and macrophages close to amyloid deposits in the brain and pancreas, respectively (Figure 8B-C). These cell types play similar roles in these tissues, and their recruitment to amyloid aggregates is consistent with prior observations [83, 84]. In the pancreas, an increased proportion of activated stellate cells can also be observed close to amyloid deposits (Figure 8C), which is in line with stellate cell activation after beta cell loss and which contributes to islet fibrosis and macrophage recruitment [85]. LPE lipids have broadly been shown to inhibit M1 macrophage polarization [86]. Interestingly, LPE 18:1 was found to be abundant in both islet niches 4 and 5 (Figures 7F and 7I-J), which are dominated by beta cells and alpha cells, respectively (Figure 7D). LPE 18:0 was most abundant in niche 4 (Figures 7F), which is dominated by beta cells, and LPE O-16:1 was most abundant in niche 3 (Figures 7F-H), the acinar pancreatic niche. This indicates that not only do LPE lipids vary in spatial distribution within the pancreas, but it also suggests that specific LPE species, such as LPE 18:1 and LPE 18:0, are more likely to be involved in macrophage polarization and recruitment. When cell lines were treated with various LPE species, LPE 18:1 induced a calcium gradient within cells, whereas 18:0 did not [87]. The differing spatial distributions of these lipids within islets suggest that these islet niches may experience differences in LPE-induced calcium transients. Differential regulation of calcium-regulatory genes was also observed near amyloid plaques in the brain and pancreas. Among these genes are CXCR4, SLIT3, S100A12, SCGN, and CXCL10 (Table S4).

Fibrosis likely also plays an important role in neurodegeneration, involving microglia, endothelial cells, and pericytes [88, 89]. We observe increased proportions of each of these cell types close to amyloid in the brain. However, it should be noted that the inclusion of CAA pathology in the amyloid mask may bias these results somewhat towards cell types associated with the vasculature, including endothelial cells and pericytes. The oxidized lipid Paz-PC was detected in the vascular region of the AD brain (Figure 2D). This lipid has been shown to promote mitochondrial permeability and inflammation, and its detection suggests a potential role in amyloid pathology in the brain [90, 91]. The KIT gene, encoding a receptor tyrosine kinase, has been linked to both AD and T2D and has been assigned a role in fibrosis [92]. Interestingly, this gene was found to have lower expression in proximity to amyloid in the brain and pancreas (Table S4).

Comparing amyloid proximity-related gene expression changes across the pancreas and brain, an overlap of 16 genes was observed (Table S4). One of these genes is SOX2, which has been linked to AD pathogenesis and plays a role in processing amyloid precursor protein (APP) [93]. Decreased levels of SOX2 have been reported in relation to AD pathology, and it may serve a neuroprotective role. In line with this, we observed decreased expression levels of SOX2 closer to amyloid deposits.

The CAV1 gene, encoding the Caveolin-1 (CAV1) membrane protein, has also been assigned a neuroprotective role in neurodegenerative diseases through its interaction with membrane lipid rafts, and here we also observe a decreased expression of CAV1 closer to amyloid in the brain (Table S4). In the pancreas, CAV1 is involved in insulin signaling and has been implicated in the development of T2D by contributing to insulin resistance, oxidative stress, and inflammation [95]. Interestingly, CAV1 has previously been put forward as a link between AD and T2D, with T2D models presenting lower brain CAV1 levels paired with increased tau hyperphosphorylation [96], and brains of T2D patients having lower CAV1 expression and higher beta amyloid levels [97].

Although we offer a significant advance in single-section and serial-section multimodal spatial workflows, some limitations still apply. Tissue deformations and tears may arise across tissue sections and between imaging platforms, and while *image2image* handles even extreme cases quite well, some local artifacts in the registration may still occur. Thus, we observed that cross-modality comparisons at the single-cell level are optimally made on the same tissue section. In our study, cell type and amyloid plaque annotations from MxIF and small molecule staining could reliably be used to mine ST profiles. We performed cross-modality mining between serial sections with larger tissue structures. For example, we interrogated IMS data using the niches derived from ST data acquired from serial tissue sections. Similarly, we performed a global correlation between the gene densities of amyloid-responsive genes and IMS ions at the whole-section level. Our approach provides multiple options for cross-modality mining of datasets from serial sections and demonstrates the power of acquiring multiple data modalities on a single tissue section.

In conclusion, our novel multimodal spatial approach enables the exploration of potential molecular markers associated with pathological hallmarks within their spatial context. The modular framework is readily generalizable to work with other omics platforms and is widely applicable to other organs and diseases.

## Resource availability

### Lead Contact

Requests for further information and resources should be directed to and will be fulfilled by the lead contact, Jeffrey M. Spraggins.

### Materials availability

This study did not generate new unique reagents.

### Data and code availability

Data have been deposited within the Human BioMolecular Atlas Program (HuBMAP) repository as HBM524.TBLD.976 and are publicly available as of the date of publication. All original code has been deposited in GitHub and is publicly available at https://github.com/vandeplaslab/image2image and https://github.com/joanagoncalveslab as of the date of publication.

## Acknowledgements

This work was supported by NIH common fund project number 1U54EY032442-01, NIH T32 DK101003, the Vanderbilt Diabetes Research and Training Center (NIH grant DK20593), NIH project number 3OT2OD033759-01S1 subaward number 1090719-473495, National Institute on Aging (NIA) (R01AG078803 awarded to JS), Human Pancreas Analysis Program DK106755, DK112217, and DK123716, and the Department of Veterans Affairs (BX000666).

## Author contributions

Conceptualization, A.R.S.K., J.M.S, and A.C.P.; Methodology, A.R.S.K, R.L., L.M.; Software, R.L., L.M., R.V.P., J.G.; Formal Analysis, R.L., L.M., A.R.S.K.; Investigation, A.R.S.K., C.F.S., C.M., M.C.M., A.E., T.P., K.A., L.V.A.; Writing – Original Draft, A.R.S.K., R.L., C.F.S., C.M.; Writing-Review & Editing, J.M.S., M.A.F., A.R.S.K., R.L., L.M., C.F.S., M.M., C.M., K.A.; Funding Acquisition, J.M.S., R.V.P., J.G., A.C.P., M.S.; Resources, J.M.S., A.C.P., M.S.

## Methods

### Human Brain Sample Preparation

The frontal cortex of a 70-year-old Caucasian male brain donor with severe Alzheimer’s disease (AD) and cerebral amyloid angiopathy (CAA) was studied. Brain tissue blocks were flash frozen in liquid nitrogen and stored at −80 °C. Tissue blocks were then embedded in 15% fish gelatin and sectioned at 10 μm thickness using a CM3050 S cryostat (Leica Biosystems, Wetzlar, Germany). This study used two serial tissue sections from this brain donor. The MALDI section was thaw-mounted onto an indium tin oxide (ITO) coated glass slide (Delta Technologies, Loveland, CO), while the section for Xenium and CODEX was mounted onto a 10X Genomics Xenium slide. Autofluorescent images were taken of the section prior to MALDI imaging with standard DAPI, eGFP, DSRed, and CY5 filters using a Zeiss AxioScan.Z1 slide scanner (Carl Zeiss Microscopy GmbH, Oberkochen, Germany), equipped with a Colibri7 LED light source.

### Human Pancreas Sample Preparation and Staining

Human pancreas tissue in the Vanderbilt Pancreas Biorepository was processed in Pittsburgh by R. Bottino in a manner including multiple processing and fixation methods, including paraformaldehyde (PFA)-fixed cryosections as described previously [98]. This study utilized two 10 μm-thick serial tissue sections from a 43-year-old black male donor with type 2 diabetes. The section used for MALDI was thaw-mounted onto an ITO-coated glass slide (Delta Technologies, Loveland, CO), while the section for Xenium and CODEX was mounted onto a 10X Genomics Xenium slide.

### MALDI IMS Tissue Preparation

The 10 μm brain tissue section was stored in a vacuum-sealed bag at −80°C for ∼3 weeks, and prior to imaging, it was brought to room temperature under vacuum for 30 minutes in the dark. Autofluorescence microscopy images were acquired on a Zeiss AxioScan.Z1 slide scanner. Three washes with 150 mM ammonium formate were performed on ice for 45 seconds each to remove salts, followed by drying with a flow of nitrogen gas at ∼3 psi. Slides were then sublimated with approximately 0.8 µg/mm^2^ with 4-Dimethylaminocinnamaldehyde (DMACA) solubilized in Tetrahydrofuran.

The human pancreas tissue section was thawed under vacuum, imaged for autofluorescence, and washed with ammonium formate as described above. Matrix was applied using an in-house developed sublimation apparatus at a concentration of 8 mg/mL DMACA in acetone. A total of 16 mg of matrix was applied to the sample surface. Heat annealing was performed to facilitate crystal formation at a temperature of 100°C for 10 seconds.

### MALDI IMS Data Acquisition

MALDI IMS data from brain tissue were acquired at 10 µm spatial resolution in negative ionization qTOF mode with a scan range of *m*/*z* 250 to 2500 on a Bruker timsTOF FleX mass spectrometer (Bruker Daltonics). Laser power was set to 18% and each pixel consisted of 25 shots from a SmartBeam 3D 10 kHz frequency tripled Nd:YAG laser (355 nm). Prior to data acquisition, a spot of red phosphorus on the target plate was used for mass calibration.

MALDI IMS data from pancreas tissue were acquired at 10 µm spatial resolution in negative ionization qTOF mode with a scan range of *m*/*z* 400 to 2000 on a Bruker timsTOF Pro mass spectrometer (Bruker Daltonics). Laser power was set to 30% and each pixel consisted of 200 shots from a SmartBeam 3D 10 kHz frequency tripled Nd:YAG laser (355 nm). Prior to data acquisition, ESI tune mix (Agilent Technologies) was used for mass calibration.

### Xenium In Situ Gene Expression

Xenium v1 *in situ* gene expression and imaging were performed per the manufacturer’s guidelines. Tissues were thaw-mounted on Xenium slides with fiducial etchings to enable Xenium analyzer recognition. The sections were then fixed and permeabilized to allow DNA padlock probes to hybridize to target RNA, followed by ligation of bound DNA probes and rolling cycle enzymatic amplification to increase the target signal. For the brain, the 266-plex pre-designed human brain panel was used (Table S5) for cell type classification and mRNA expression of genes associated with AD pathologies. For the pancreas, the 377-plex Human Multi-Tissue and Cancer panel was used (Table S5) to target various cell types of the pancreas. Prior to imaging on the Xenium Analyzer, slides were quenched of autofluorescence and stained with DAPI for cell segmentation. Cyclical fluorescent probe hybridization, imaging, decoding, and data processing were all performed onboard the instrument (instrument software version 1.9.2.0, analysis version xenium-1.9.0.0). Immediately following the completion of the Xenium runs, the slides were removed from the Xenium slide cassettes and kept in 1x PBS at 4°C until the post-Xenium PhenoCycler was performed as described below.

### Post-Xenium PhenoCycler Multiplexed Immunofluorescence

Antibodies for PhenoCycler were purchased pre-conjugated from Akoya Biosciences or procured from other vendors and conjugated internally using the Akoya Biosciences PhenoCycler Conjugation Kit. Pancreas tissue was stained using an established antibody panel (Table S6). Brain tissue was stained using a combination of custom-conjugated and commercially available antibodies (Akoya Biosciences) (Table S6). Briefly, tissue sections were fixed with ice cold acetone for 10 minutes, washed twice with hydration buffer, fixed using 1.6% PFA in hydration buffer for 10 minutes, washed with hydration buffer twice, followed by staining buffer for 20 minutes, then stained with antibody cocktail solution in blocking buffer for three hours at room temperature. Tissue was washed with staining buffer and post-staining fixation was performed in 1.6% PFA in staining buffer for 10 minutes. Tissues were washed three times in DPBS, followed by ice-cold methanol for 5 minutes. Tissues were washed with DPBS three times followed by final fixation in 2% Fixative Reagent for 20 minutes at room temperature. Tissues were stored in storage buffer until imaging. A reporter plate was prepared according to the manufacturer’s specifications, and cyclical imaging was performed using an Akoya PhenoCycler Fusion.

### Amyloid staining

After PhenoCycler imaging, flow cells were removed from each slide by soaking in histoclear for approximately 16 hours. Brain sections were fixed in 4% paraformaldehyde (PFA) and then photobleached for 48 hours. Sections were then stained for β-amyloid using Thiazine Red (1µM, Cat# 2150-33-6, Chemsavers) along with DAPI Fluoromount. Pancreas sections were fixed with 4% PFA, washed using PBST, and stained with Thioflavin S for 10 minutes, washed three times in PBS, and coverslipped with DAPI Fluoromount as previously reported [99]. Imaging was performed using a Zeiss AxioScan.Z1 with a 10x objective.

### Xenium data analysis

Following the Xenium runs, the counts data were exported from the Xenium Analyzer for processing and analysis with R (version 4.4.0) [100] and RStudio (version 2024.04.2) [101], using the Seurat package (version 5.0.3) [102] as a basis. For tool parameters not specified, the defaults were used. Cells with 0 transcript counts were filtered out and the data were visually inspected. The counts were then normalized using *SCTransform* [103], after which, first a principal component analysis was performed, followed by uniform manifold approximation and projection (UMAP) dimensionality reduction on the first 30 principal components (PCs) for plotting [104]. The first 30 PCs were also used for clustering with *FindNeighbors* and *FindClusters* (using the Leiden algorithm with resolution = 0.2 and method = “igraph”) [105]. as a basis. For tool parameters not specified, the defaults were used. Cells with 0 transcript counts were filtered out and the data was visually inspected. The counts were then normalized using *SCTransform* [103], after which first a principal component analysis was performed, followed by uniform manifold approximation and projection (UMAP) dimensionality reduction on the first 30 principal components (PCs) for plotting [104]. The first 30 PCs were also used for clustering with *FindNeighbors* and *FindClusters* (using the Leiden algorithm with resolution = 0.2 and method = “igraph”) [105].

Raw counts were used for cell type prediction with RCTD [106], as implemented in the spacexr package (version 2.2.1) [107]. Publicly available datasets were used as reference for cell type predictions; a scRNA-seq dataset generated by Baron *et al*. [49] for the pancreas and a snRNA-seq dataset from the Allen Institute for the brain [108]. The raw counts of the pancreas reference dataset were used and the count matrices of the 4 human donors from that study were merged into a combined reference. For the brain reference dataset, the raw exonic counts per gene were used, only cells with a sub class label were included, and this is also the label that was used for the cell type predictions. One gene of the Xenium Human Brain Expression panel (*CEMIP2*) was present in the reference dataset under its previous name (*TMEM2*) and so this gene name was updated to have matching gene names in the panel and the reference data. RCTD objects were created with UMI_min = 50, UMI_min_sigma = 150 and CELL_MIN_INSTANCE = 5. RCTD was then run in doublet mode and the first cell type in the results was used to assign cell types in Xenium data. Using the assigned cell types, a niche analysis was performed using the *BuildNicheAssay* function in Seurat, with neighbors.k = 100 and niches.k = 6 for the brain and niches.k = 5 for the pancreas.

### Integration of spatial transcriptomics and histopathology

After registration of the Xenium spatial transcriptomics and histopathology modalities to the same coordinate system, these were integrated using Python (version 3.10.14). The transformed cell segmentation coordinates were grouped by cell into shapely (version 2.1.0) polygons and combined into a geopandas (version 1.0.1) GeoDataFrame. Invalid cell polygon geometries were made valid with the *make_valid* function of shapely. The GeoDataFrame was then parsed for use with the SpatialData framework (version 0.2.2). For the amyloid masks, the registered GeoJSON files from QuPath could directly be read and parsed into a SpatialData-compatible GeoDataFrame using the *ShapesModel.parse* function. For each cell its distance to the nearest amyloid pathological feature was determined by first finding the closest point of the amyloid mask with the *nearest_points* shapely function, and then the distance to that point was determined with the *distance* function. Cell overlap with amyloid directly, or with 20- and 40-micron dilation masks, was determined with the *overlaps* function. The distance and overlap cell annotations were combined with the cell segmentation and amyloid masks, as well as the registered Xenium DAPI and Thiazine Red or Thioflavin S staining images into a single SpatialData object to allow for combined visualizations.

The amyloid distance and overlap annotations were added to the processed transcriptomics data for further analysis in R. Cells were binned to determine cell type proportions at different distances from amyloid. Clustering per cell type was performed to identify subpopulations, potentially associated with amyloid proximity. For this the data was subset and analyzed per cell type, following the same procedure as for the full data, but with clustering using the first 10 PCs and a resolution of 0.1. For each cell type, genes showing expression differences associated with proximity to amyloid were identified by fitting a linear model with limma (version 3.60.0) [109], using voom-transformed counts (with TMM effective library size normalization ) as the outcome and distance to amyloid as the predictor. Significant associations were defined as those with *p* < 0.05 after Bonferroni multiple testing correction. Due to heavily skewed cell densities across the amyloid distances, this analysis was limited to cells within a distance of 100 μm from amyloid, where the cell density was more uniform.

### MALDI IMS Data Preprocessing

MALDI IMS data were exported from the Bruker timsTOF file format (.d) to a custom binary format. Each pixel/frame contained centroid peaks spanning the entire acquisition range. These were reconstructed into a pseudo-profile mass spectrum using Bruker Daltonics’ TDF-SDK (v2.21). The data were m/z-aligned using at least six peaks commonly present in the majority of pixels, utilizing the msalign library (v0.2.0) [110, 111]. The mass axis of each dataset was calibrated using a minimum of four theoretical masses to achieve an accuracy of approximately ±1 ppm.

Subsequently, the MALDI IMS data were normalized using a total ion current (TIC) approach, and an average mass spectrum was computed based on all pixels. This average spectrum underwent peak-picking, resulting in the detection of 1,420 peaks in brain samples and 889 peaks in pancreas samples. It is important to note that isotopic peaks were not removed prior to proceeding with the unsupervised and classification workflows. Each peak list was then used to extract ion centroid data, which was integrated within a window of ± approximately 5 ppm. This value could vary slightly depending on the bin spacing in the specific mass spectrum.

To facilitate cell type and niche analysis, Xenium-, PhenoCycler-, and stain-based masks were transformed to align with the IMS coordinate system. For each mask, an average mass spectrum was extracted, providing detailed insights for further analysis.

### Image Co-registration and Multimodal Alignment

The microscopy registration process was performed in two steps. First, we co-registered all modalities within the same section using the *elastix* framework integrated into the *image2image* application. [48]. For slide 1, this included the Xenium DAPI stain maximum intensity projection (MIP) for the ST modality, PhenoCycler DAPI stain for the MxIF modality, autofluorescence microscopy (AF), and thioflavin S/thiazine red stains for histopathology; and for slide 2 this included both pre-IMS AF and post-IMS AF for the IMS modality. For this step, we used *rigid* and *affine* transformations, as substantial differences (e.g., deformations, tears, or other artefacts) are not expected within the same tissue section. Registration nodes were established between imaging modalities to ensure optimal registration alignment (Figure S2). For example, in the registration of the brain sample (Figure S2A), the Xenium DAPI stain, PhenoCycler, AF, and thiazine red stains were co-registered. Specifically, the PhenoCycler modality was registered to the AF image, and the Xenium DAPI image was registered to the PhenoCycler DAPI image. The transformations obtained during the registration of PhenoCycler to AF were then applied to the Xenium modality, thereby aligning all three modalities into the same coordinate system. This essentially enabled an indirect registration of the Xenium transcript locations to the AF image.

Due to variation between serial sections, we used *rigid, affine,* and *B-spline* transformations to perform non-linear registration. For example, the PhenoCycler modality from slide 1 was registered to the pre-IMS AF image from slide 2 (Figure S2B). Similar procedures were employed for other serial sections, such as the pancreas, where the thioflavin S stain image (slide 1) was aligned with the pre-IMS AF image (slide 2) (Figure S2B). Following the completion of these registration steps, a visual inspection of all images was performed at both tissue and cellular scales to ensure accuracy (Figure S3). Furthermore, the MALDI IMS datasets were manually registered to the post-IMS autofluorescence microscopy images, which display the desorption marks created by the laser during acquisition. This manual registration was performed within the *image2image* GUI, where 8-12 fiducial markers were selected in both modalities to estimate the *affine* transformation.

### Supervised Machine Learning and Shapley Additive Explanations

Masks were utilized to develop an eXtreme Gradient Boosting (XGBoost) classification model, designed to identify predicted cell types or niches. We adopted a one-versus-all approach for multi-class classification, where each classification task aims to differentiate the positive class against all other pixels in the dataset. To identify which ion species have marker-like relationships to each mask, we employed Shapley Additive Explanations (SHAP). SHAP quantifies the importance of each m/z species in recognizing the mask within the classification model, providing both experiment-wide (global) and per-pixel (local) importance scores.

Global SHAP importance scores rank all m/z (or gene) species by decreasing relevance for recognizing a specific class, highlighting a set of highly discriminative molecular species that represent potential biomarker candidates. Conversely, local SHAP importance scores assess the direction of relevance (positive or negative monotonic correlation) and evaluate the significance of the relationship between an ion species’ intensity and a pixel’s likelihood of belonging to a class.

The classification models were trained using a 67%/33% train/test split and implemented with the scikit-learn (v1.3.0) [112], XGBoost (v2.1.1) [113], and SHAP (0.46.0) [52, 114] libraries in Python (v3.9.18). Given that some masks cover less than 5% of the dataset, leading to extremely unbalanced data, we used imbalanced-learn (v0.11.0) to adjust the dataset balance by over-sampling the minority class with synthetic data augmentation and under-sampling the majority class through random subsampling.

Hyper-parameter optimization for each mask was performed using a grid-search approach on a subset of the training data to maximize accuracy. The best-performing parameters were then applied to construct the classification models. SHAP was subsequently used to quantify the importance of each ion species for the classification tasks, calculating a global SHAP score by averaging the magnitudes of the local SHAP scores across all pixels.

In the case of cellular niches, a one-verse-rest binary classification was performed at the pixel level for each niche. SHAP values were computed using the TreeSHAP algorithm, which efficiently and precisely quantifies the contribution of each gene or lipid to the model’s prediction [53]. Each feature was assigned both a global and local SHAP importance score. The global SHAP score is obtained by calculating the mean of the magnitude of the local SHAP importance scores across all pixels, with higher values indicating greater importance for niche classification.

We ranked the SHAP scores in descending order of importance, creating a shortlist of molecular species that may serve as useful biomarker candidates. In the summary bubble plots, the size of each marker represents the global SHAP importance score for each ion (column) for a given classification task (row). Additionally, we determined whether a biomarker candidate is positively or negatively correlated with a given ion species by calculating the Spearman rank-order correlation coefficient (*p*) between the mean-centered intensity and the Shapley values of each molecular ion. The Spearman correlation coefficient ranges from −1 to 1, where magnitudes greater than 0.2 are considered significant. In the bar charts and summary bubble plots, the marker color corresponds to this correlation coefficient for each ion species, illustrating the strength and direction of their relationships.

### Segmentation of microscopy images

Segmentation of PhenoCycler and amyloid staining images was performed in QuPath v.0.5.1. Amyloid plaque segmentation masks were generated from thiazine red (brain) and thioflavin S (pancreas) stains using thresholding and positive cell selection tools with the parameters below:

| *Thiazine Red Thresholding Parameters* |  | *Thioflavin S Thresholding Parameters* |  |
| --- | --- | --- | --- |
| **Resolution** | 1.30 $\mu\text{m}/\text{px}$ | **Resolution** | 1 $\mu\text{m}/\text{px}$ |
| **Channel** | DsRed | **Channel** | EGFP |
| **Prefilter** | Gaussian | **Prefilter** | Gaussian |
| **Smoothing Sigma** | 3 | **Smoothing Sigma** | 2 |
| **Above 4000** | Amyloid plaque | **Above 4500** | Amyloid |
| **Below 4000** | Unclassified | **Below 4500** | Unclassified |
| **Region** | Everywhere | **Region** | Everywhere |

## References

1. Seferbekova, Z., et al., Spatial biology of cancer evolution. Nat Rev Genet, 2023. 24(5): p. 295–313.

2. Hu, T., et al., Single-cell spatial metabolomics with cell-type specific protein profiling for tissue systems biology. Nat Commun, 2023. 14(1): p. 8260.

3. Hu, B.C., The human body at cellular resolution: the NIH Human Biomolecular Atlas Program. Nature, 2019. 574(7777): p. 187–192.

4. Gulati, G.S., et al., Profiling cell identity and tissue architecture with single-cell and spatial transcriptomics. Nat Rev Mol Cell Biol, 2025. 26(1): p. 11–31.

5. Jain, S. and M.T. Eadon, Spatial transcriptomics in health and disease. Nat Rev Nephrol, 2024. 20(10): p. 659–671.

6. Jain, S., et al., Advances and prospects for the Human BioMolecular Atlas Program (HuBMAP). Nat Cell Biol, 2023. 25(8): p. 1089–1100.

7. Colley, M.E., et al., High-Specificity Imaging Mass Spectrometry. Annu Rev Anal Chem (Palo Alto Calif), 2024. 17(1): p. 1–24.

8. Marshall, C.R., et al., Untangling Alzheimer’s disease with spatial multi-omics: a brief review. Front Aging Neurosci, 2023. 15: p. 1150512.

9. Djambazova, K.V., et al., Resolving the Complexity of Spatial Lipidomics Using MALDI TIMS Imaging Mass Spectrometry. Anal Chem, 2020. 92(19): p. 13290–13297.

10. Neumann, E.K., et al., Multimodal Imaging Mass Spectrometry: Next Generation Molecular Mapping in Biology and Medicine. J Am Soc Mass Spectrom, 2020. 31(12): p. 2401–2415.

11. Kruse, A.R.S. and J.M. Spraggins, Uncovering Molecular Heterogeneity in the Kidney With Spatially Targeted Mass Spectrometry. Front Physiol, 2022. 13: p. 837773.

12. Gale, S.A., D. Acar, and K.R. Daffner, Dementia. Am J Med, 2018. 131(10): p. 1161–1169.

13. Long, J.M. and D.M. Holtzman, Alzheimer Disease: An Update on Pathobiology and Treatment Strategies. Cell, 2019. 179(2): p. 312–339.

14. Attems, J. and K.A. Jellinger, The overlap between vascular disease and Alzheimer’s disease--lessons from pathology. BMC Med, 2014. 12: p. 206.

15. Biffi, A. and S.M. Greenberg, Cerebral Amyloid Angiopathy: A Systematic Review. J Clin Neurol, 2011. 7(1): p. 1–9.

16. Boyle, P.A., et al., Cerebral amyloid angiopathy and cognitive outcomes in community-based older persons. Neurology, 2015. 85(22): p. 1930–6.

17. Yamada, M. and H. Naiki, Cerebral amyloid angiopathy. Prog Mol Biol Transl Sci, 2012. 107: p. 41–78.

18. Gatti, L., et al., Understanding the Pathophysiology of Cerebral Amyloid Angiopathy. Int J Mol Sci, 2020. 21(10).

19. Greenberg, S.M., et al., Cerebral amyloid angiopathy and Alzheimer disease - one peptide, two pathways. Nat Rev Neurol, 2020. 16(1): p. 30–42.

20. Solopova, E., et al., *Fatal iatrogenic cerebral* β*-amyloid-related arteritis in a woman treated with lecanemab for Alzheimer’s disease*. Nat Commun, 2023. 14(1): p. 8220.

21. Sun, H., et al., IDF Diabetes Atlas: Global, regional and country-level diabetes prevalence estimates for 2021 and projections for 2045. Diabetes Res Clin Pract, 2022. 183: p. 109119.

22. Federation, I.D., *IDF Diabetes Atlas*. 2025: Brussels, Belgium.

23. O’Brien, T.D., et al., Islet amyloid polypeptide: a review of its biology and potential roles in the pathogenesis of diabetes mellitus. Vet Pathol, 1993. 30(4): p. 317–32.

24. Westermark, P., A. Andersson, and G.T. Westermark, Islet amyloid polypeptide, islet amyloid, and diabetes mellitus. Physiol Rev, 2011. 91(3): p. 795–826.

25. Jha, D., et al., Spatial neurolipidomics-MALDI mass spectrometry imaging of lipids in brain pathologies. J Mass Spectrom, 2024. 59(3): p. e5008.

26. Prentice, B.M., et al., Imaging mass spectrometry enables molecular profiling of mouse and human pancreatic tissue. Diabetologia, 2019. 62(6): p. 1036–1047.

27. Fantini, J. and N. Yahi, Molecular insights into amyloid regulation by membrane cholesterol and sphingolipids: common mechanisms in neurodegenerative diseases. Expert Rev Mol Med, 2010. 12: p. e27.

28. García-Viñuales, S., et al., *The interplay between lipid and A*β *amyloid homeostasis in Alzheimer’s Disease: risk factors and therapeutic opportunities*. Chem Phys Lipids, 2021. 236: p. 105072.

29. Imai, Y., et al., Connecting pancreatic islet lipid metabolism with insulin secretion and the development of type 2 diabetes. Ann N Y Acad Sci, 2020. 1461(1): p. 53–72.

30. Prentki, M., B.E. Corkey, and S.R.M. Madiraju, Lipid-associated metabolic signalling networks in pancreatic beta cell function. Diabetologia, 2020. 63(1): p. 10–20.

31. Schnackenberg, L.K., et al., MALDI imaging mass spectrometry: an emerging tool in neurology. Metab Brain Dis, 2022. 37(1): p. 105–121.

32. Fernández-Calle, R., et al., APOE in the bullseye of neurodegenerative diseases: impact of the APOE genotype in Alzheimer’s disease pathology and brain diseases. Mol Neurodegener, 2022. 17(1): p. 62.

33. Caprioli, R.M., T.B. Farmer, and J. Gile, Molecular imaging of biological samples: localization of peptides and proteins using MALDI-TOF MS. Anal Chem, 1997. 69(23): p. 4751–60.

34. Angel, P.M. and R.M. Caprioli, Matrix-assisted laser desorption ionization imaging mass spectrometry: in situ molecular mapping. Biochemistry, 2013. 52(22): p. 3818–28.

35. Norris, J.L. and R.M. Caprioli, Analysis of tissue specimens by matrix-assisted laser desorption/ionization imaging mass spectrometry in biological and clinical research. Chem Rev, 2013. 113(4): p. 2309–42.

36. Kaya, I., et al., Histology-Compatible MALDI Mass Spectrometry Based Imaging of Neuronal Lipids for Subsequent Immunofluorescent Staining. Anal Chem, 2017. 89(8): p. 4685–4694.

37. Angel, P.M., et al., MALDI Imaging Mass Spectrometry of N-glycans and Tryptic Peptides from the Same Formalin-Fixed, Paraffin-Embedded Tissue Section. Methods Mol Biol, 2018. 1788: p. 225–241.

38. Angel, P.M., K. Norris-Caneda, and R.R. Drake, In Situ Imaging of Tryptic Peptides by MALDI Imaging Mass Spectrometry Using Fresh-Frozen or Formalin-Fixed, Paraffin-Embedded Tissue. Curr Protoc Protein Sci, 2018. 94(1): p. e65.

39. Esselman, A.B., et al., In situ molecular profiles of glomerular cells by integrated imaging mass spectrometry and multiplexed immunofluorescence microscopy. Kidney Int, 2025. 107(2): p. 332–337.

40. Marx, V., Method of the Year: spatially resolved transcriptomics. Nat Methods, 2021. 18(1): p. 9–14.

41. Janesick, A., et al., High resolution mapping of the tumor microenvironment using integrated single-cell, spatial and in situ analysis. Nat Commun, 2023. 14(1): p. 8353.

42. Black, S., et al., CODEX multiplexed tissue imaging with DNA-conjugated antibodies. Nat Protoc, 2021. 16(8): p. 3802–3835.

43. Patterson, N.H., et al., Advanced Registration and Analysis of MALDI Imaging Mass Spectrometry Measurements through Autofluorescence Microscopy. Analytical Chemistry, 2018. 90(21): p. 12395–12403.

44. Nathan Heath, P., et al., Autofluorescence microscopy as a label-free tool for renal histology and glomerular segmentation. bioRxiv, 2021: p. 2021.07.16.452703.

45. Sud, M., et al., LMSD: LIPID MAPS structure database. Nucleic Acids Res, 2007. 35(Database issue): p. D527–32.

46. Conroy, M.J., et al., LIPID MAPS: update to databases and tools for the lipidomics community. Nucleic Acids Res, 2024. 52(D1): p. D1677–d1682.

47. Migas, L.G., Image2image. retrieved from https://github.com/vandeplaslab/image2image-docs?tab=readme-ov-file, 2023.

48. Klein, S., et al., *elastix: a toolbox for intensity-based medical image registration*. IEEE Trans Med Imaging, 2010. 29(1): p. 196–205.

49. Baron, M., et al., A Single-Cell Transcriptomic Map of the Human and Mouse Pancreas Reveals Inter- and Intra-cell Population Structure. Cell Syst, 2016. 3(4): p. 346–360.e4.

50. Preston, A.N., D.A. Cervasio, and S.T. Laughlin, Visualizing the brain’s astrocytes. Methods Enzymol, 2019. 622: p. 129–151.

51. Rajasekaran, S.A., et al., *Na,*K-ATPase subunits as markers for epithelial-mesenchymal transition in cancer and fibrosis. Mol Cancer Ther, 2010. 9(6): p. 1515–24.

52. Tideman, L.E.M., et al., Automated biomarker candidate discovery in imaging mass spectrometry data through spatially localized Shapley additive explanations. Anal Chim Acta, 2021. 1177: p. 338522.

53. Lundberg, S.M.E., G.; Lee, S., *Consistent Individualized Feature Attribution for Tree Ensembles.* 2019.

54. Lehrer, S. and P.H. Rheinstein, RORB, an Alzheimer’s disease susceptibility gene, is associated with viral encephalitis, an Alzheimer’s disease risk factor. Clin Neurol Neurosurg, 2023. 233: p. 107984.

55. Hijazi, S., A.B. Smit, and R.E. van Kesteren, Fast-spiking parvalbumin-positive interneurons in brain physiology and Alzheimer’s disease. Mol Psychiatry, 2023. 28(12): p. 4954–4967.

56. Saade, M., et al., The Role of GPNMB in Inflammation. Front Immunol, 2021. 12: p. 674739.

57. Bonham, L.W., et al., CXCR4 involvement in neurodegenerative diseases. Transl Psychiatry, 2018. 8(1): p. 73.

58. Yang, C.F., et al., Loss of GPNMB Causes Autosomal-Recessive Amyloidosis Cutis Dyschromica in Humans. Am J Hum Genet, 2018. 102(2): p. 219–232.

59. Liu, Z., et al., *CX3CR1 in microglia regulates brain amyloid deposition through selective protofibrillar amyloid-*β *phagocytosis*. J Neurosci, 2010. 30(50): p. 17091–101.

60. Lee, S., et al., CX3CR1 deficiency alters microglial activation and reduces beta-amyloid deposition in two Alzheimer’s disease mouse models. Am J Pathol, 2010. 177(5): p. 2549–62.

61. Tanno, T., et al., Slit3 regulates cell motility through Rac/Cdc42 activation in lipopolysaccharide-stimulated macrophages. FEBS Lett, 2007. 581(5): p. 1022–6.

62. Dulyaninova, N.G., et al., S100A4 regulates macrophage invasion by distinct myosin-dependent and myosin-independent mechanisms. Mol Biol Cell, 2018. 29(5): p. 632–642.

63. Ambartsumian, N., J. Klingelhöfer, and M. Grigorian, The Multifaceted S100A4 Protein in Cancer and Inflammation. Methods Mol Biol, 2019. 1929: p. 339–365.

64. Jiang, X., et al., Temporal expression patterns of insulin-like growth factor binding protein-4 in the embryonic and postnatal rat brain. BMC Neurosci, 2013. 14: p. 132.

65. Iwadate, H., et al., Actions of insulin-like growth factor binding protein-5 (IGFBP-5) are potentially regulated by tissue kallikrein in rat brains. Life Sci, 2003. 73(24): p. 3149–58.

66. Son, J.W., et al., Glia-Like Cells from Late-Passage Human MSCs Protect Against Ischemic Stroke Through IGFBP-4. Mol Neurobiol, 2019. 56(11): p. 7617–7630.

67. Zhang, W., et al., Decorin is a pivotal effector in the extracellular matrix and tumour microenvironment. Oncotarget, 2018. 9(4): p. 5480–5491.

68. D’Amico M, A., et al., Biological function and clinical relevance of chromogranin A and derived peptides. Endocr Connect, 2014. 3(2): p. R45–54.

69. Remes, S.M., et al., PCSK2 expression in neuroendocrine tumors points to a midgut, pulmonary, or pheochromocytoma-paraganglioma origin. Apmis, 2020. 128(11): p. 563–572.

70. Mulvihill, K.G., Presynaptic regulation of dopamine release: Role of the DAT and VMAT2 transporters. Neurochem Int, 2019. 122: p. 94–105.

71. Chen, M.K., et al., VMAT2 and dopamine neuron loss in a primate model of Parkinson’s disease. J Neurochem, 2008. 105(1): p. 78–90.

72. Rege, T.A. and J.S. Hagood, Thy-1 as a regulator of cell-cell and cell-matrix interactions in axon regeneration, apoptosis, adhesion, migration, cancer, and fibrosis. Faseb j, 2006. 20(8): p. 1045–54.

73. Meijer, B., R.B. Gearry, and A.S. Day, The role of S100A12 as a systemic marker of inflammation. Int J Inflam, 2012. 2012: p. 907078.

74. Madácsy, T., P. Pallagi, and J. Maleth, Cystic Fibrosis of the Pancreas: The Role of CFTR Channel in the Regulation of Intracellular Ca(2+) Signaling and Mitochondrial Function in the Exocrine Pancreas. Front Physiol, 2018. 9: p. 1585.

75. Krüger, C., et al., AQP8 is a crucial H(2)O(2) transporter in insulin-producing RINm5F cells. Redox Biol, 2021. 43: p. 101962.

76. Wu, X., et al., Unveiling the role of CXCL10 in pancreatic cancer progression: A novel prognostic indicator. Open Med (Wars), 2025. 20(1): p. 20241117.

77. Burgener, R., et al., Purification and characterization of a major phosphatidylserine-binding phosphoprotein from human platelets. Biochem J, 1990. 269(3): p. 729–34.

78. McMahon, K.A., et al., Identification of intracellular cavin target proteins reveals cavin-PP1alpha interactions regulate apoptosis. Nat Commun, 2019. 10(1): p. 3279.

79. Kaya, I., et al., Spatial lipidomics reveals brain region-specific changes of sulfatides in an experimental MPTP Parkinson’s disease primate model. NPJ Parkinsons Dis, 2023. 9(1): p. 118.

80. Kaya, I., et al., Brain region-specific amyloid plaque-associated myelin lipid loss, APOE deposition and disruption of the myelin sheath in familial Alzheimer’s disease mice. J Neurochem, 2020. 154(1): p. 84–98.

81. Kaya, I., et al., Shedding Light on the Molecular Pathology of Amyloid Plaques in Transgenic Alzheimer’s Disease Mice Using Multimodal MALDI Imaging Mass Spectrometry. ACS Chem Neurosci, 2018. 9(7): p. 1802–1817.

82. He, X., et al., Deregulation of sphingolipid metabolism in Alzheimer’s disease. Neurobiol Aging, 2010. 31(3): p. 398–408.

83. Kamata, K., et al., *Islet amyloid with macrophage migration correlates with augmented* β*-cell deficits in type 2 diabetic patients*. Amyloid, 2014. 21(3): p. 191–201.

84. Mallach, A., et al., Microglia-astrocyte crosstalk in the amyloid plaque niche of an Alzheimer’s disease mouse model, as revealed by spatial transcriptomics. Cell Rep, 2024. 43(6): p. 114216.

85. Yang, Y., et al., *Pancreatic stellate cells in the islets as a novel target to preserve the pancreatic* β*-cell mass and function*. J Diabetes Investig, 2020. 11(2): p. 268–280.

86. Park, S.J. and D.S. Im, 2-Arachidonyl-lysophosphatidylethanolamine Induces Anti-Inflammatory Effects on Macrophages and in Carrageenan-Induced Paw Edema. Int J Mol Sci, 2021. 22(9).

87. Makiyama, F., et al., Differential effects of structurally different lysophosphatidylethanolamine species on proliferation and differentiation in pre-osteoblast MC3T3-E1 cells. Sci Rep, 2025. 15(1): p. 466.

88. D’Ambrosi, N. and S. Apolloni, Fibrotic Scar in Neurodegenerative Diseases. Front Immunol, 2020. 11: p. 1394.

89. Laredo, F., J. Plebanski, and A. Tedeschi, Pericytes: Problems and Promises for CNS Repair. Front Cell Neurosci, 2019. 13: p. 546.

90. Lidman, M., et al., The oxidized phospholipid PazePC promotes permeabilization of mitochondrial membranes by Bax. Biochim Biophys Acta, 2016. 1858(6): p. 1288–97.

91. Bagchi, A.K., et al., IL-10 attenuates OxPCs-mediated lipid metabolic responses in ischemia reperfusion injury. Sci Rep, 2020. 10(1): p. 12120.

92. Martinez-Anton, A., et al., KIT as a therapeutic target for non-oncological diseases. Pharmacol Ther, 2019. 197: p. 11–37.

93. Sarlak, G. and B. Vincent, The Roles of the Stem Cell-Controlling Sox2 Transcription Factor: from Neuroectoderm Development to Alzheimer’s Disease? Mol Neurobiol, 2016. 53(3): p. 1679–1698.

94. Wang, H.K., et al., Hyperglycemia compromises the ischemia-provoked dedifferentiation of cerebral pericytes through p21-SOX2 signaling in high-fat diet-induced murine model. Diab Vasc Dis Res, 2021. 18(1): p. 1479164121990641.

95. Tang, W., et al., Caveolin-1, a novel player in cognitive decline. Neurosci Biobehav Rev, 2021. 129: p. 95–106.

96. Wu, J., et al., High glucose induces formation of tau hyperphosphorylation via Cav-1-mTOR pathway: A potential molecular mechanism for diabetes-induced cognitive dysfunction. Oncotarget, 2017. 8(25): p. 40843–40856.

97. Bonds, J.A., et al., Depletion of Caveolin-1 in Type 2 Diabetes Model Induces Alzheimer’s Disease Pathology Precursors. J Neurosci, 2019. 39(43): p. 8576–8583.

98. Brissova, M., et al., α *Cell Function and Gene Expression Are Compromised in Type 1 Diabetes*. Cell Rep, 2018. 22(10): p. 2667–2676.

99. Walker, J.T., et al., Genetic risk converges on regulatory networks mediating early type 2 diabetes. Nature, 2023. 624(7992): p. 621–629.

100. Team, R.C., R: A Language and Environment for Statistical Computing. 2022.

101. Team, R., RStudio: Integrated Development Environment for R. 2020.

102. Hao, Y., et al., Dictionary learning for integrative, multimodal and scalable single-cell analysis. Nat Biotechnol, 2024. 42(2): p. 293–304.

103. Choudhary, S. and R. Satija, Comparison and evaluation of statistical error models for scRNA-seq. Genome Biol, 2022. 23(1): p. 27.

104. Melville, L.M.J.H.J., UMAP: Uniform Manifold Approximation and Projection for Dimension Reduction. arXiv, 2020. 1802.03426.

105. Traag, V.A., L. Waltman, and N.J. van Eck, From Louvain to Leiden: guaranteeing well-connected communities. Sci Rep, 2019. 9(1): p. 5233.

106. Cable, D.M., et al., Robust decomposition of cell type mixtures in spatial transcriptomics. Nat Biotechnol, 2022. 40(4): p. 517–526.

107. Cable, D.M., spacexr: SpatialeXpressionR: Cell type identification and cell type-specific differential expression in spatial transcriptomics. 2024.

108. Jorstad, N.L., et al., Transcriptomic cytoarchitecture reveals principles of human neocortex organization. Science, 2023. 382(6667): p. eadf6812.

109. Ritchie, M.E., et al., *limma powers differential expression analyses for RNA-sequencing and microarray studies*. Nucleic Acids Res, 2015. 43(7): p. e47.

110. Monchamp, P., et al., Signal processing methods for mass spectrometry. Systems Bioinformatics: An Engineering Case-Based Approach, 2007: p. 101–124.

111. Migas, L.G., msalign: Spectral alignment based on MATLAB’s ‘msalign’ function. 2024.

112. Pedregosa, F.a.V. G. and Gramfort, A. and Michel, V., et al., Scikit-learn: Machine Learning in Python. Journal of Machine Learning Research, 2011. 12: p. 2825–2830.

113. Guestrin, T.C.a.C., XGBoost: A Scalable Tree Boosting System. CoRR, 2016. abs/1603.02754(1603.02754).

114. Lundberg, S.M., et al., From Local Explanations to Global Understanding with Explainable AI for Trees. Nat Mach Intell, 2020. 2(1): p. 56–67.

