## Supplemental Information for "Integrative Spatial Omics for Systems-Level Mapping of Pathological Niches"

**Supplementary Material**

**Supplementary tables (Tables S2, S3, S5, S6 provided as separate files):**

**Table S1**. Selected lipid annotations

**Table S2-** Intensity of IMS ions within unsupervised K-means clusters in the brain and pancreas.

**Table S3**- Differentially expressed genes near amyloid plaques in the brain and pancreas.

**Table S4**. Genes associated with amyloid proximity in both the brain and pancreas

**Table S5**- Xenium panels

**Table S6**- CODEX panels

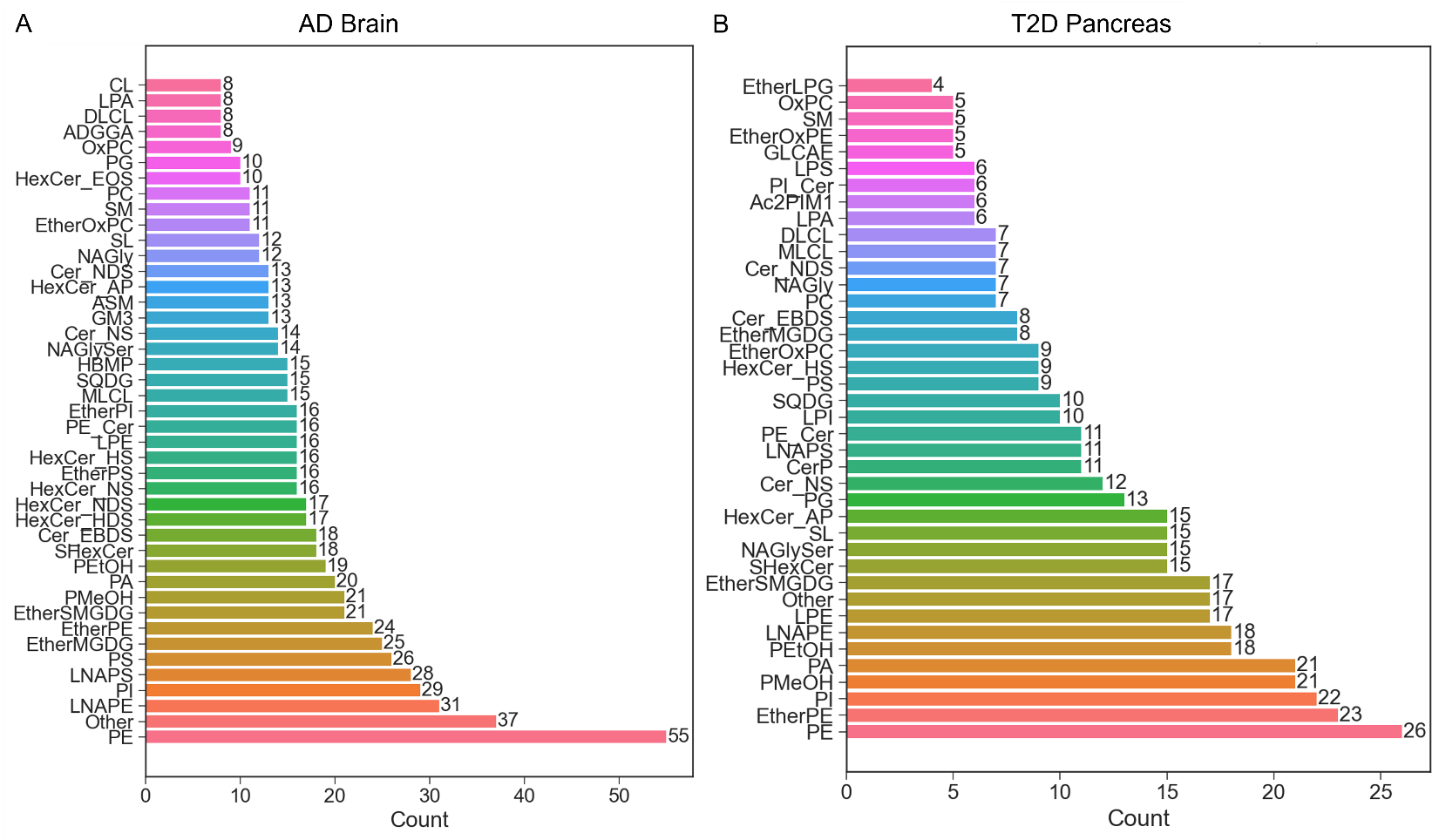

**Figure S1**. Summary of IMS lipid class annotations in AD brain (A) and T2D pancreas (B). Annotations were based on mass accuracy using the LipidMaps structural database.

**Table S1**. Selected lipid annotations

| **IMS *m/z*** | **Adduct** | **Annotation** | **Organ** |
| --- | --- | --- | --- |
| 478.295 | M-H | LPE 18:1 | brain |
| 528.31 | M-H | LPE 22:4 | brain |
| 718.539 | M-H | PE 34:0 | brain |
| 747.497 | M-H | PA 40:6 | brain |
| 766.539 | M-H | PE 38:4 | brain |
| 790.539 | M-H | PE 40:6 | brain |
| 794.569 | M-H | PE 40:4 | brain |
| 824.662 | M-H | HexCer 42:2;O3 | brain |
| 890.636 | M-H | SHexCer 42:1;O2 | brain |
| 892.617 | M+DMACA-H | SMd34:1 | brain |
| 664.416 | M-H | PaZ-PC | brain |
| 701.509 | M-H | PA 36:1 \| PA 18:0_18:1 | brain |
| 1135.806 | M-H | GA2 t38:1 | brain |
| 920.685 | M-H | SHexCer 44:0;O2 | brain |
| 904.618 | M-H | SHexCer 42:2;3O | brain |
| 747.497 | M-H | PA 40:6 \| PA 18:0_22:6 | brain |
| 892.653 | M+DMACA-H | SM d34:1 | brain |
| 788.54 | M-H | PS 36:1 \| PS 18:0_18:1 | brain |
| 885.549 | M-H | PI 38:4 \| PI 18:1_20:3 | brain |
| 885.549 | M-H | PI 38:4 \| PI 18:0_20:4 | brain |
| 799.529 | M-H | PA 44:8 | brain |
| 616.47 | M-H | CerP 34:1;O2 | pancreas |
| 670.517 | M-H | CerP 38:2;O2 | pancreas |
| 809.518 | M-H | PI 32:0 | pancreas |
| 906.34 | M-H | SHexCer 42:1;2O | pancreas |
| 833.519 | M-H | PI 34:2 | pancreas |
| 1261.816 | M-H | GM3 42:2;2O | pancreas |
| 768.554 | M-H | PE 38:3 | pancreas |
| 770.57 | M-H | PE 38:2 | pancreas |
| 478.293 | M-H | LPE 18:1 | pancreas |
| 436.283 | M-H | LPE O-16:1 | pancreas |
| 480.3 | M-H | LPE 18:0 | pancreas |
| 673.48 | M-H | PA 34:1 | pancreas |
| 647.466 | M-H | PA 32:0 | pancreas |
| 812.54 | M-H | PS 38:3 | pancreas |
| 673.481 | M-H | PA 34:1 | pancreas |
| 642.487 | M-H | CerP 36:2;O2 | pancreas |
| 462.298 | M-H | LPE O-18:2 | pancreas |
| 496.263 | M-H | LPS 16:0 | pancreas |
| 560.412 | M-H | CerP 30:1;O2 | pancreas |
| 742.54 | M-H | PE 36:2 | pancreas |
| 812.544 | M-H | PS 38:3 | pancreas |
| 885.55 | M-H | PI 38:4 | pancreas |
| 911.565 | M-H | PI 40:5 | pancreas |

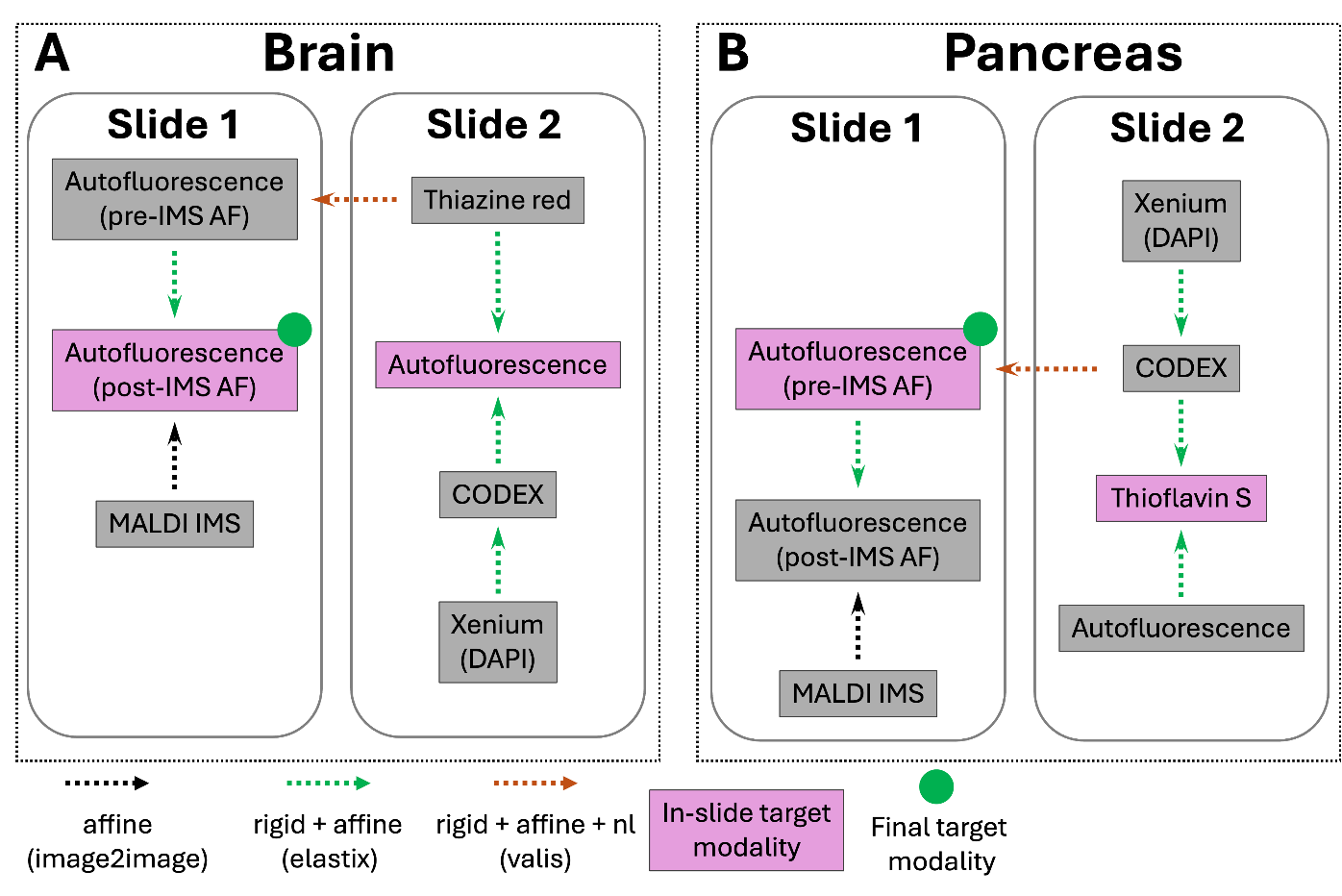

**Figure S2**. Image co-registration scheme.

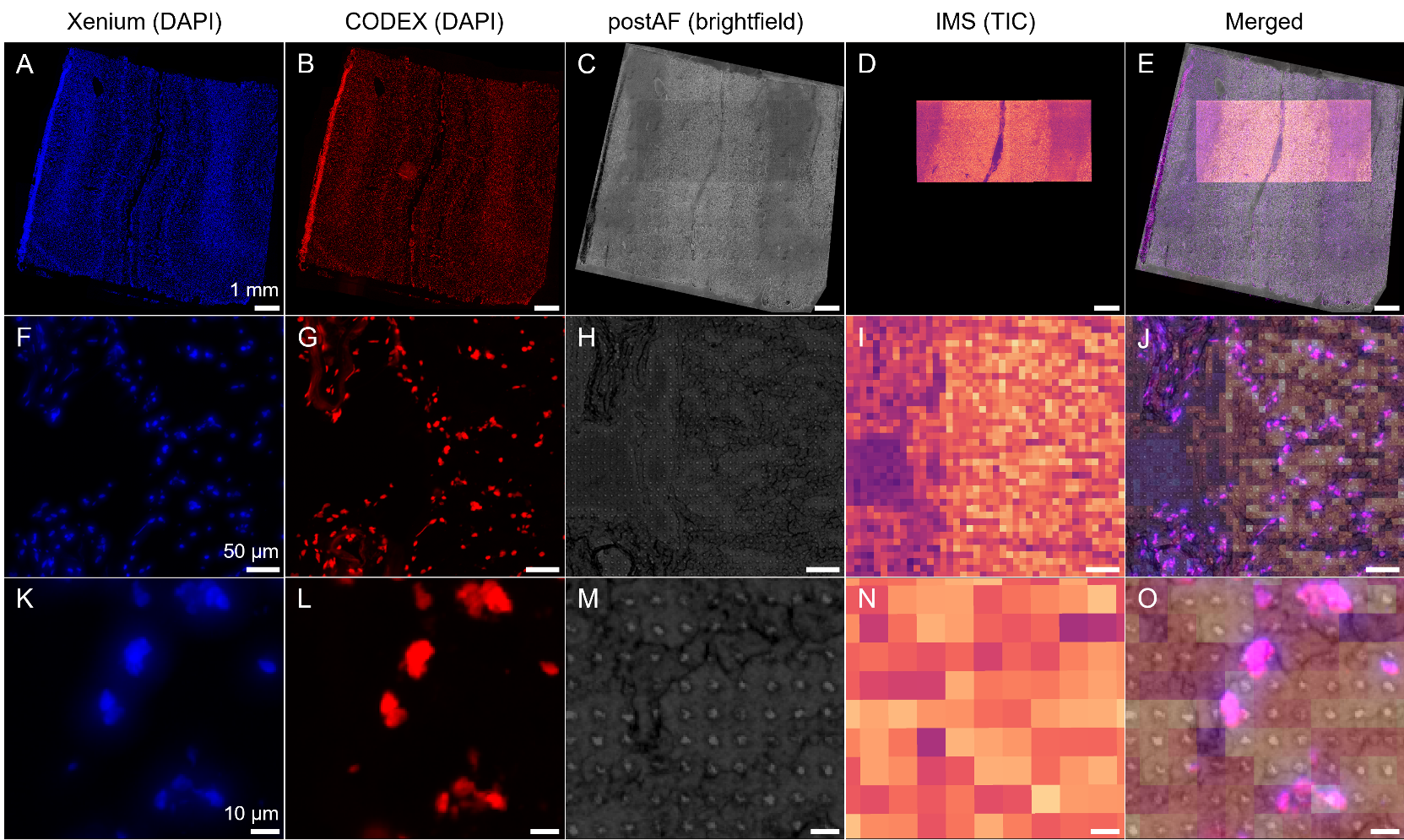

**Figure S3**. Co-registration result demonstrated in brain.

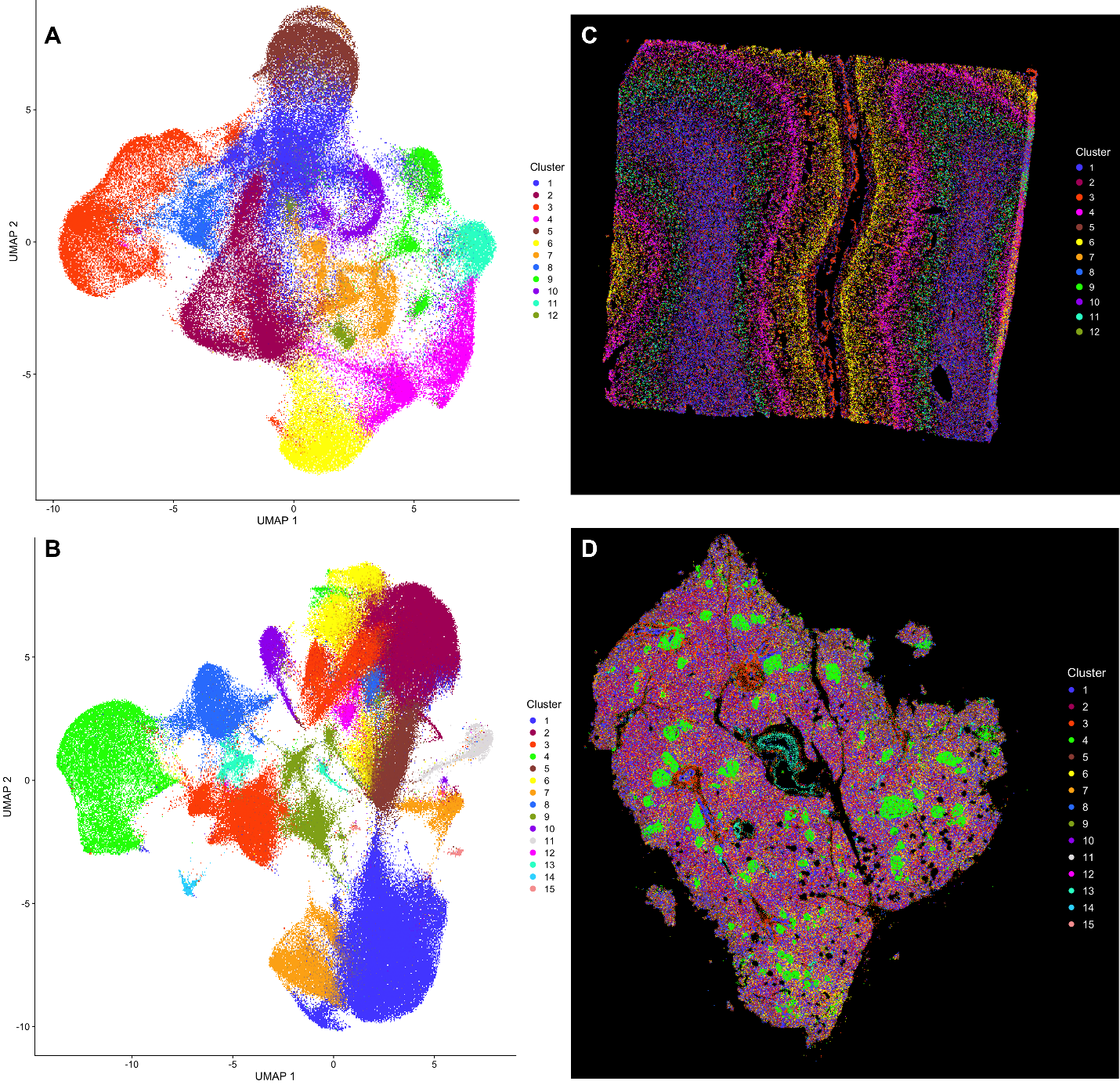

**Figure S4**. Unsupervised clustering of Xenium data in AD brain and T2D pancreas. A-B. Uniform Manifold Approximation and Projection (UMAP) was used to perform dimensionality reduction of the data in the brain (A) and pancreas (B). C-D. K-means clustering was performed to partition data in the brain (C) and pancreas (D).

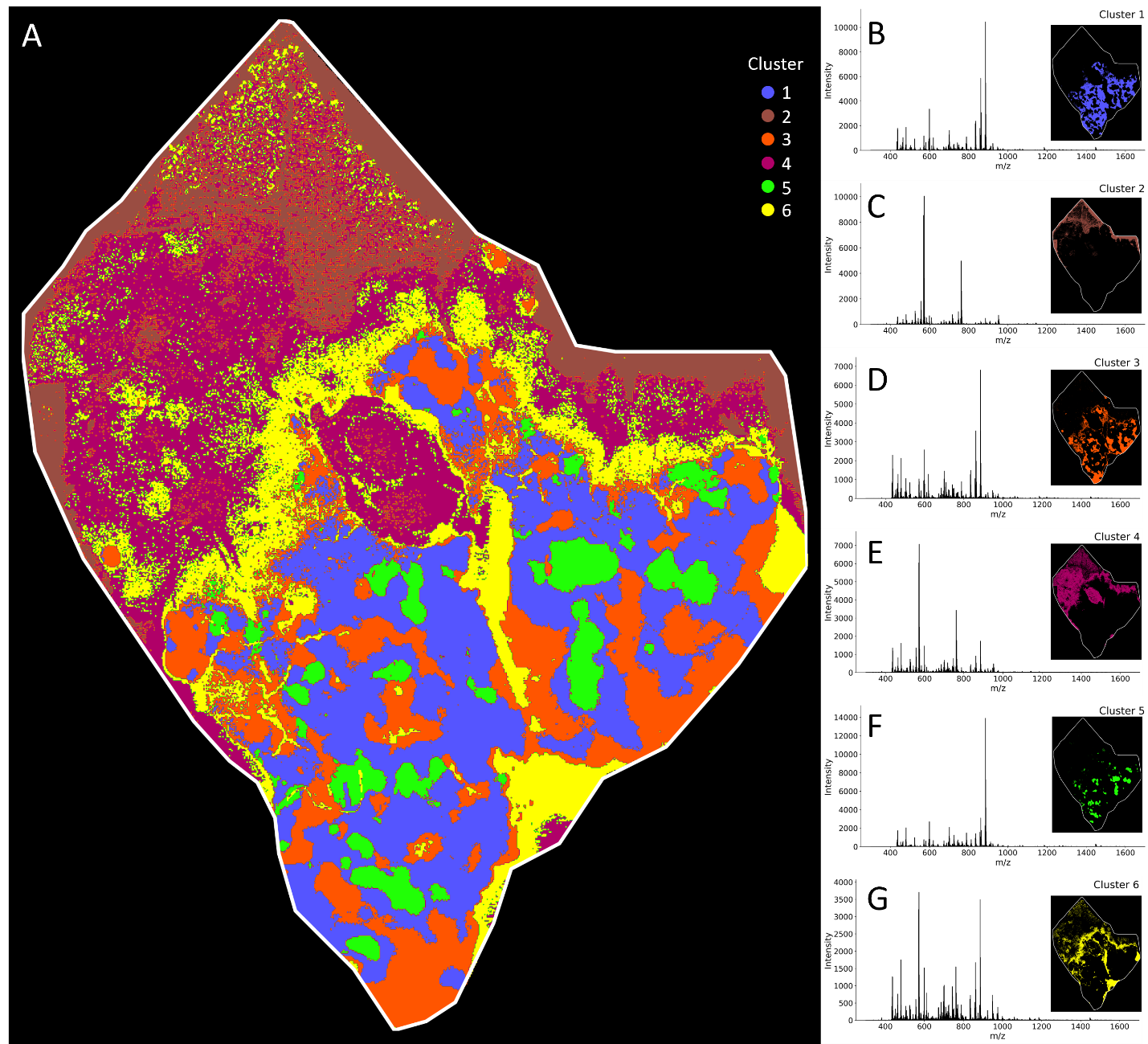

**Figure S5**. Unsupervised clustering of MALDI IMS data in T2D-pancreas. Kmeans clustering was performed to segregate MALDI IMS data into 6 clusters. Cluster identities are spatially visualized as an overlay (A) and as individual cluster images associated with an average MALDI IMS spectral profile (B-G).

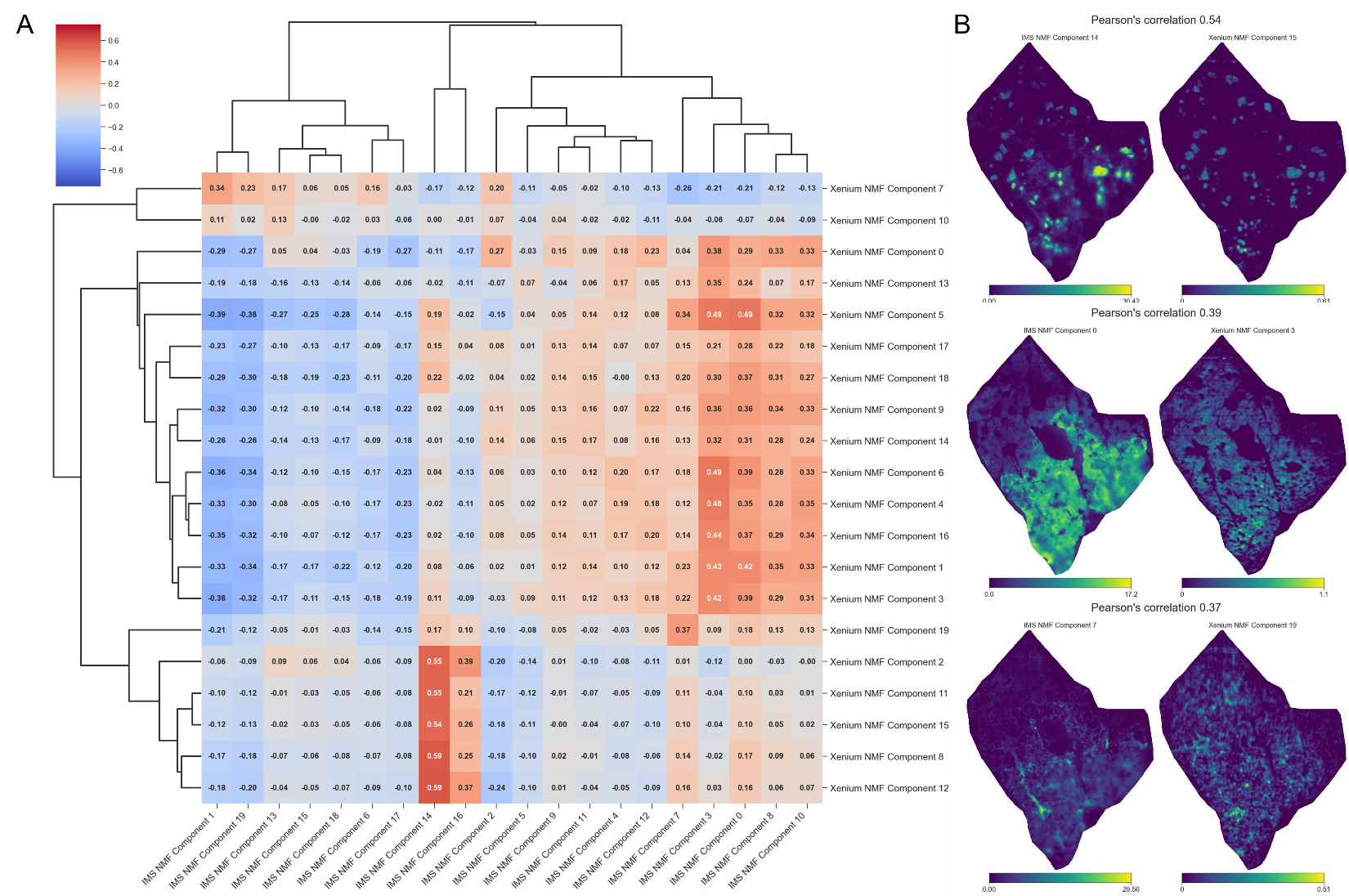

**Figure S6**. A. Pearson’s correlation of unsupervised clusters from Xenium and IMS datasets. B. Comparison of selected correlated components.

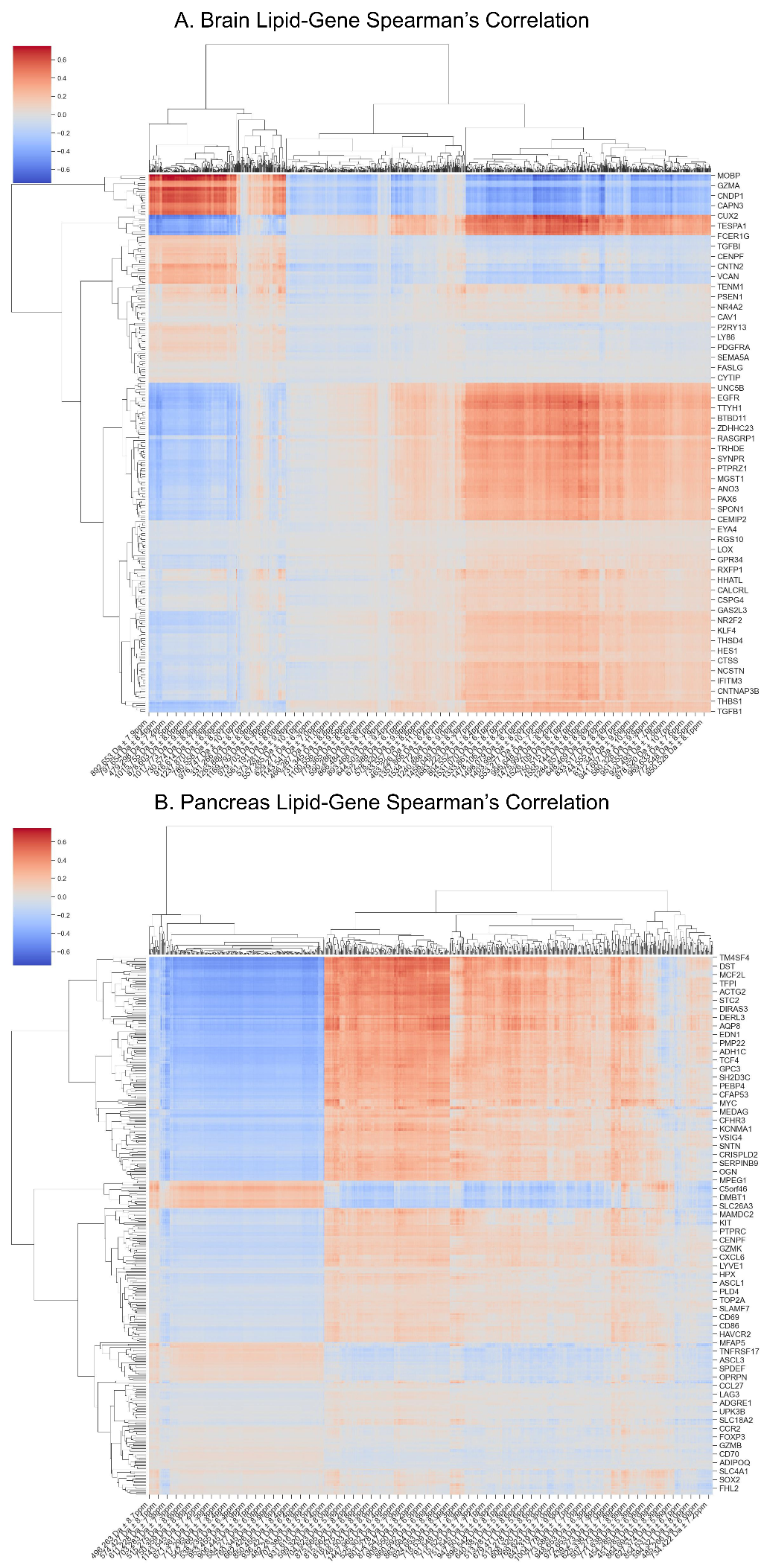

**Figure S7**. Spatial correlation of detected genes detected with Xenium and lipid ions detected with IMS in the AD brain (A) and T2D pancreas (B).

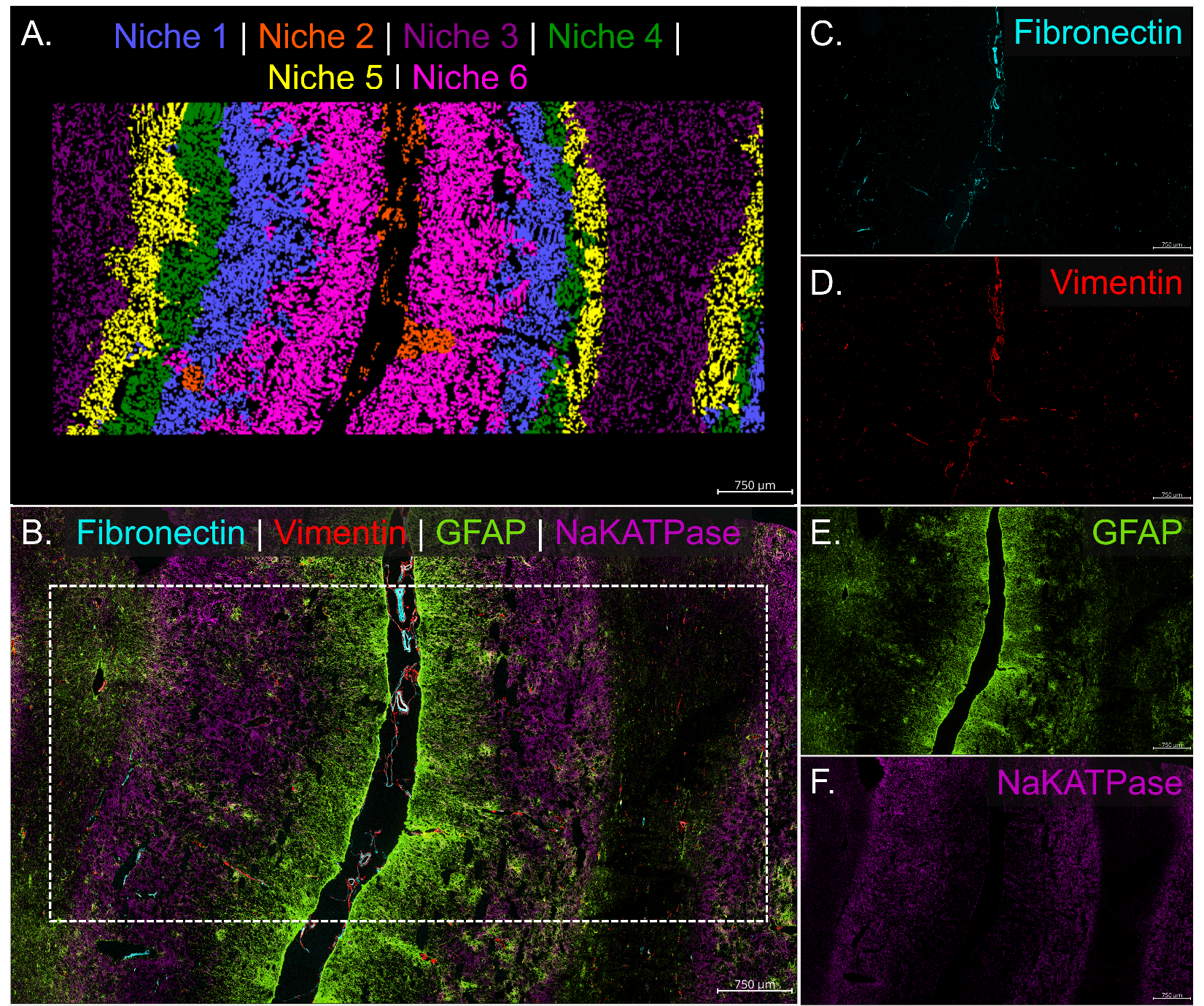

**Figure S8**. Unsupervised clustering of MALDI IMS data in AD-brain. Kmeans clustering was performed to segregate MALDI IMS data into 8 clusters. Cluster identities are spatially visualized as an overlay (A) and as individual cluster images associated with an average MALDI IMS spectral profile (B-I).

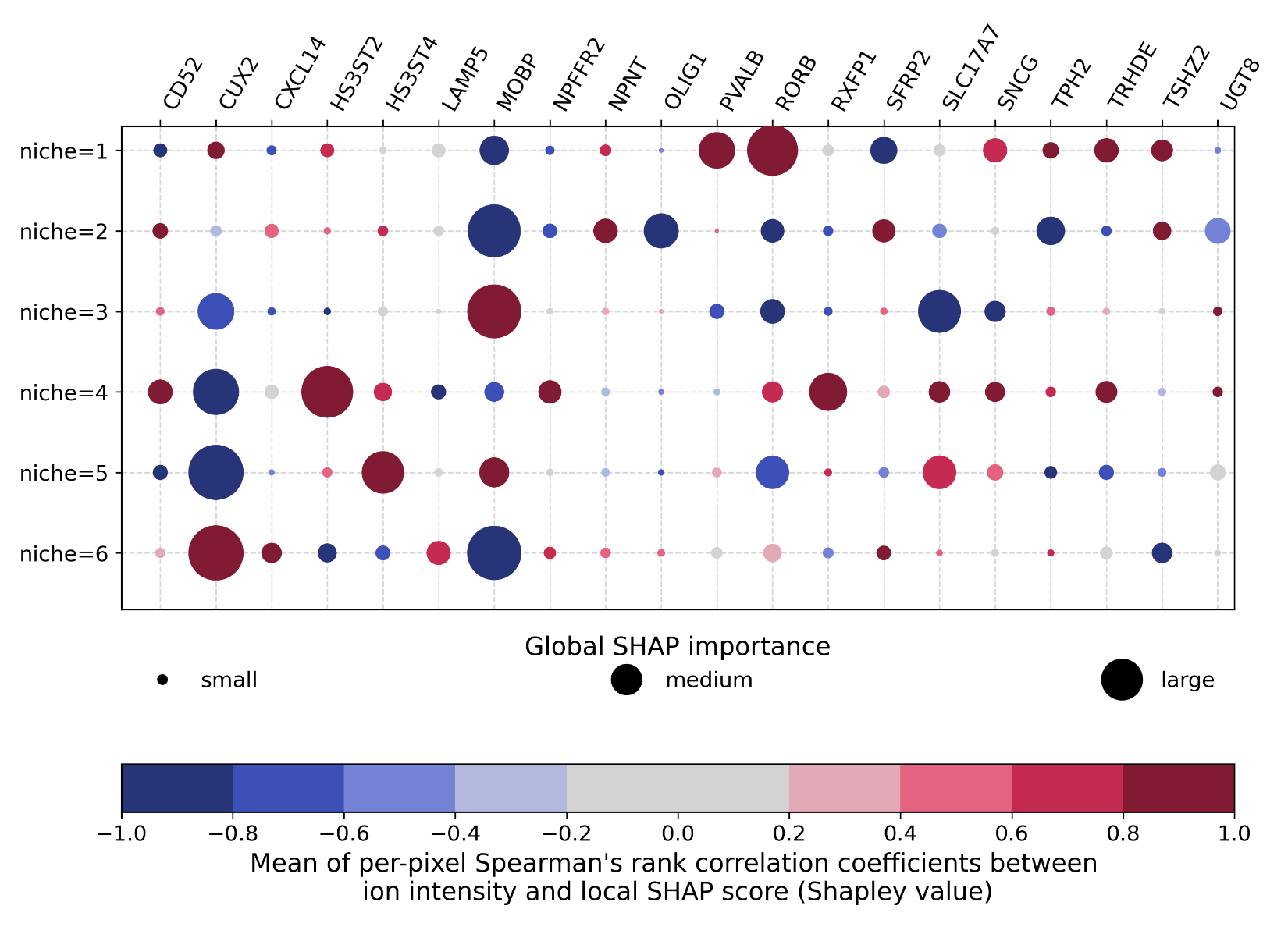

**Figure S9**. Bubble plot showing the 20 genes with the highest importance score for niche prediction in AD-brain.

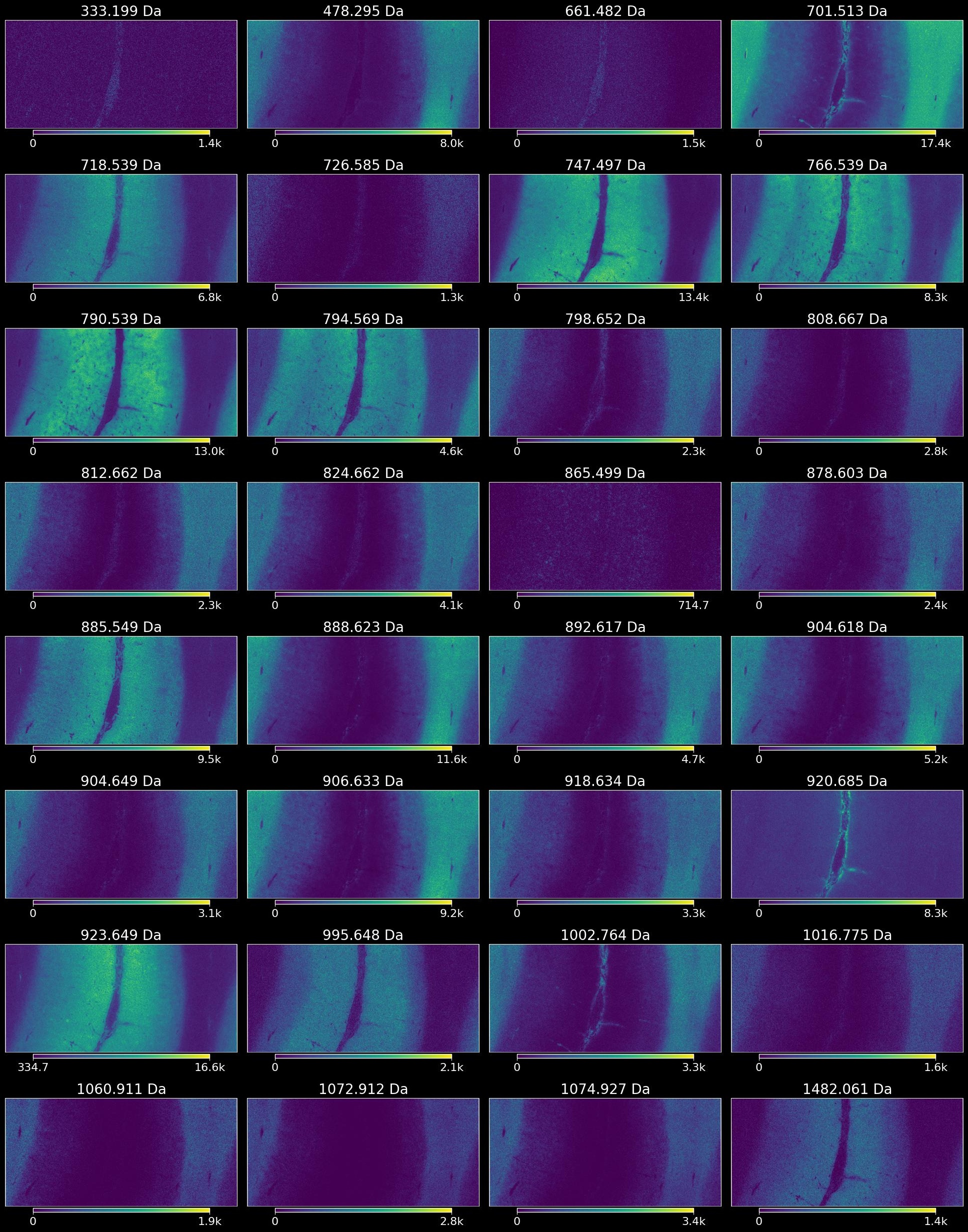

**Figure S10.** Ion images of IMS ions with the highest SHAP importance scores for prediction of niches in the AD brain.

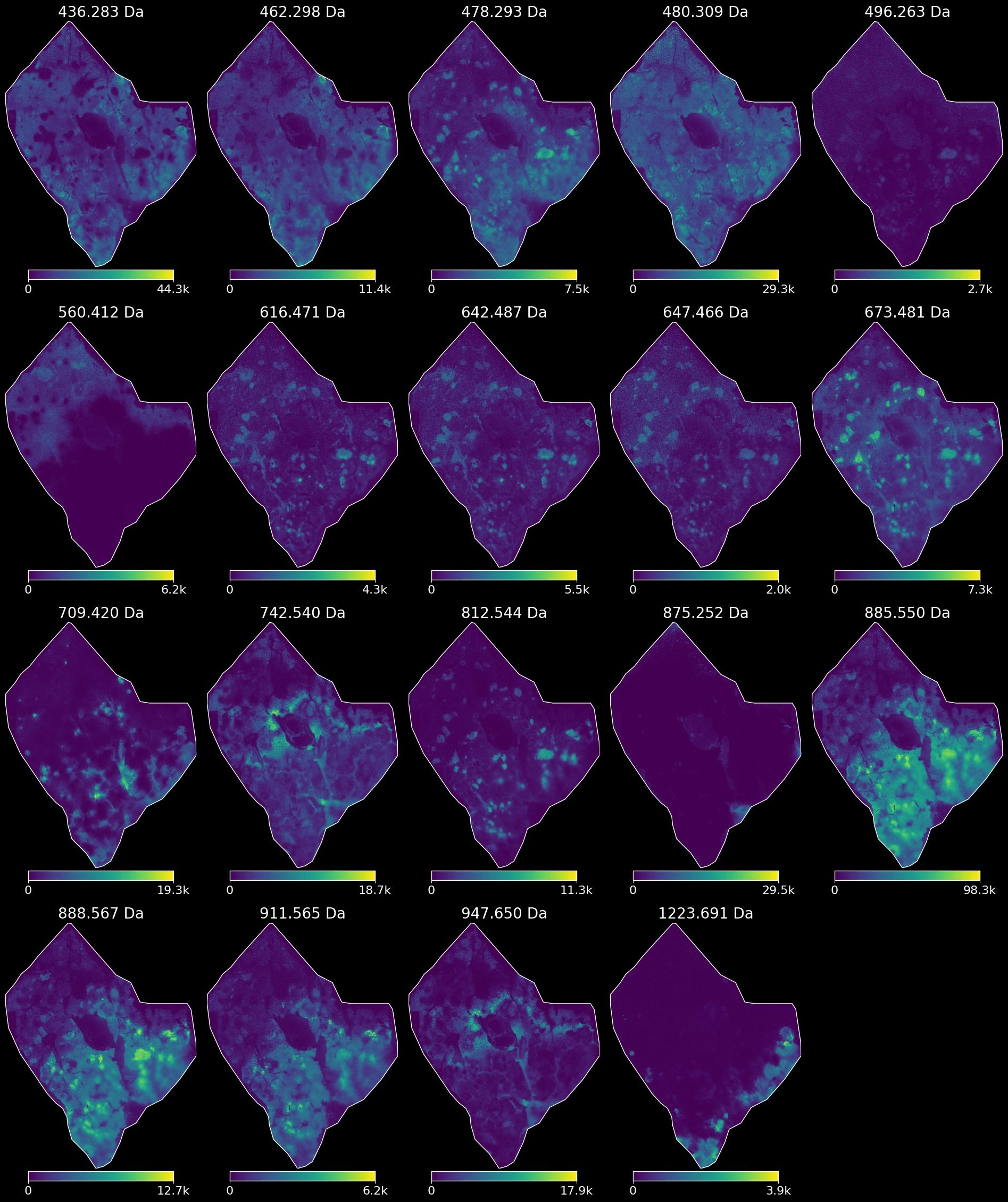

**Figure S11**. Ion images of IMS ions with the highest SHAP importance scores for prediction of niches in the T2D pancreas.

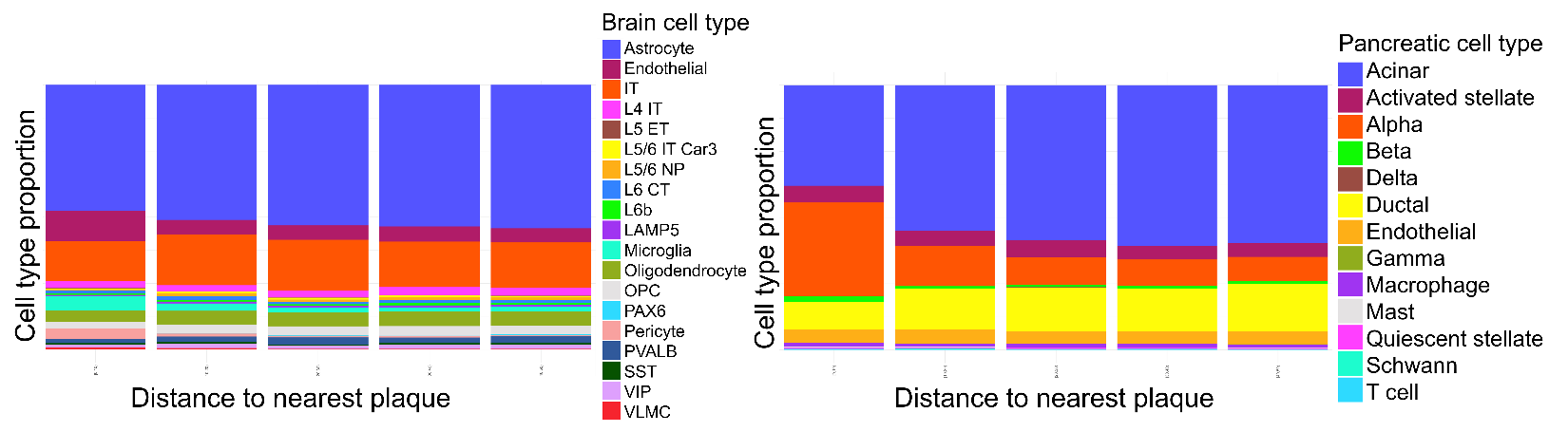

**Figure S12**. Cell type proportion as distance to plaque increases in the AD brain (left) and T2D pancreas (right).

**Table S4**. Genes associated with amyloid proximity in both the brain and pancreas

| Gene | celltype_brain | celltype_pancreas |
| --- | --- | --- |
| EGFR | Astrocyte, Endothelial | Acinar |
| SOX2 | Astrocyte, Microglia | Alpha, Acinar |
| MS4A6A | Astrocyte | Alpha, Acinar |
| FBLN1 | Astrocyte | Acinar |
| KIT | Astrocyte | Alpha, Acinar, Activated stellate |
| CXCR4 | Astrocyte, Microglia | Alpha, Acinar |
| SPI1 | Astrocyte, Endothelial | Alpha, Acinar |
| PVALB | IT | Alpha, Acinar |
| CAV1 | IT | Alpha, Acinar |
| PDGFRA | IT | Alpha, Acinar |
| CD86 | IT | Alpha, Acinar, Activated stellate |
| CD83 | Endothelial | Alpha, Acinar, Activated stellate |
| PECAM1 | Oligodendrocyte | Alpha, Acinar, Activated stellate |
| CCL5 | Oligodendrocyte | Alpha, Acinar |
| FCGR3A | Microglia | Alpha, Acinar, Activated stellate |
| SNCG | Microglia | Alpha, Acinar |

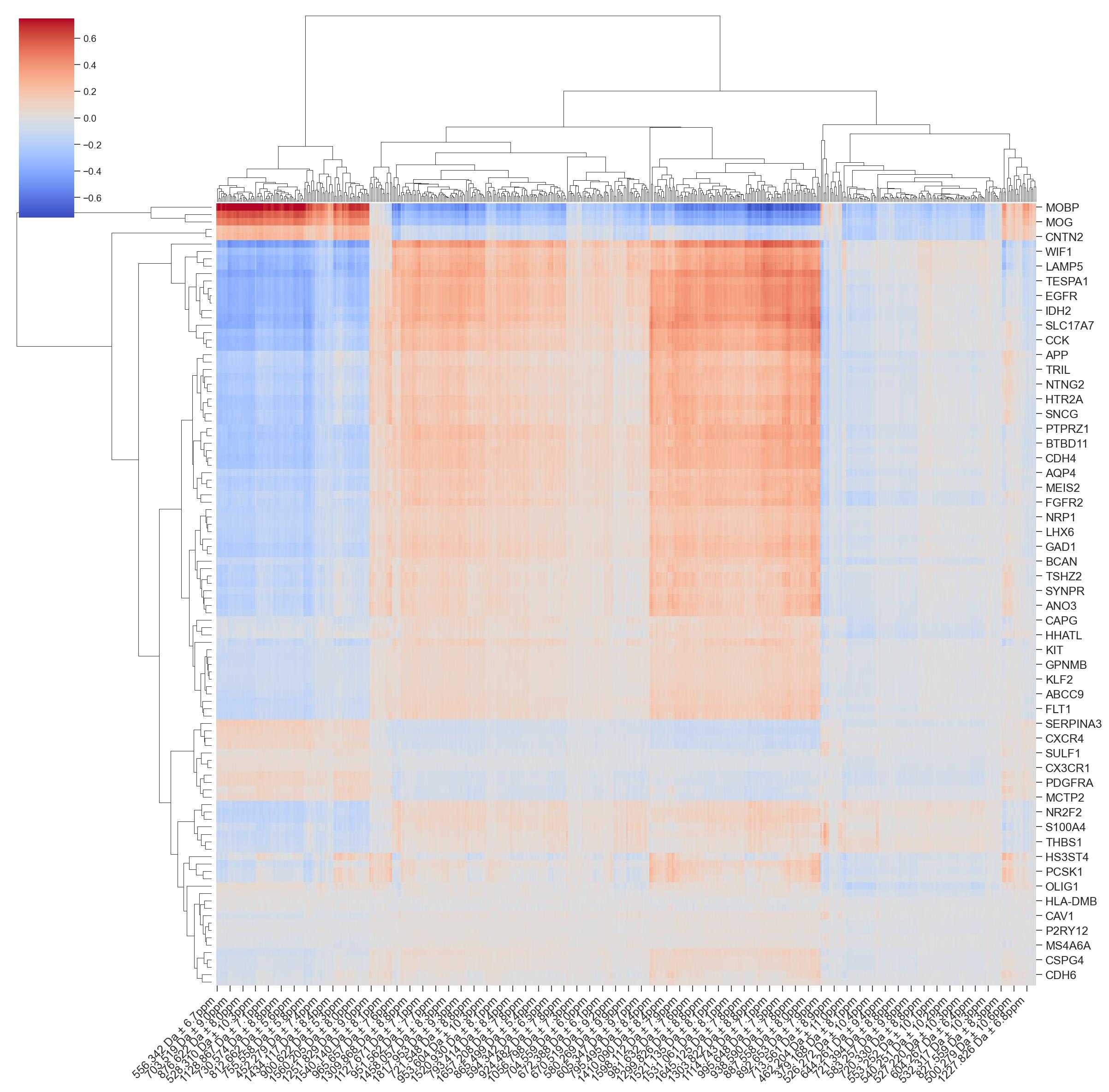

**Figure S13**. Heatmap displaying Pearson's correlation of genes differentially expressed within 100 μm of an amyloid plaque MALDI IMS lipids in the AD brain.

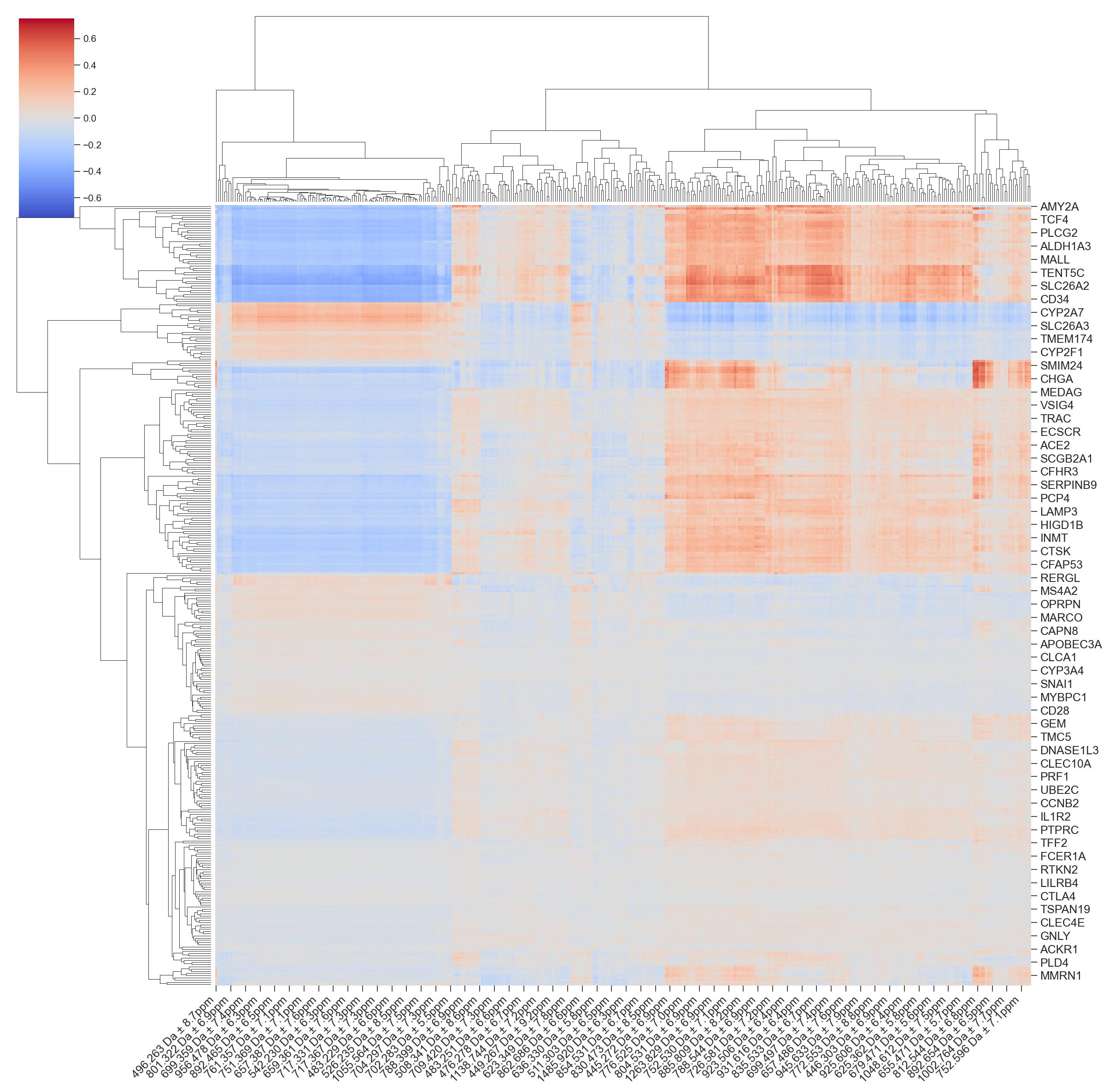

**Figure S14**. Heatmap displaying Pearson's correlation of genes differentially expressed within 100 μm of an amyloid plaque MALDI IMS lipids in the T2D pancreas.
